# Biotic homogenisation in bird communities leads to large-scale changes in species associations

**DOI:** 10.1101/2020.11.13.380956

**Authors:** Stanislas Rigal, Vincent Devictor, Pierre Gaüzère, Sonia Kéfi, Jukka T Forsman, MIra H Kajanus, Mikko Mönkkönen, Vasilis Dakos

## Abstract

**Aim:** The impact of global change on biodiversity is commonly assessed in terms of changes in species distributions, community richness and community composition. Whether and how much associations between species, *i.e*. the degree of correlation in their spatial co-occurrence, are also changing is much less documented and mostly limited to local studies of ecological networks. In this study, we quantify changes in large-scale patterns of species associations in bird communities in relation to changes in species composition.

**Location:** France.

**Time period:** 2001-2017.

**Major taxa studied:** Common breeding birds.

**Methods:** We use network approaches to build three community-aggregated indices reflecting complementary aspects of species association networks. We characterise the spatio-temporal dynamics of these indices using a large-scale and high-resolution dataset of bird co-abundances of 109 species monitored for 17 years (2001-2017) from 1,969 sites across France. We finally test whether spatial and temporal changes in species association networks are related to species homogenisation estimated as the spatio-temporal dynamics of β-diversity and the proportion of habitat generalists. The consistency of these relationships is tested across three main habitats, namely woodland, grassland and human settlements.

**Results:** We document a directional change in association-based indices in response to modifications in β-diversity and in the proportion of generalists in space and time. Weaker associations and sparser networks were related to lower β-diversity and a higher proportion of generalists, suggesting an overlooked aspect of biotic homogenisation affecting species associations. We report that this overall pattern is not constant across habitats, with opposite relationships between biotic homogenisation and change in species association networks in urban versus forest communities suggesting distinct homogenisation processes.

**Main Conclusions:** Although species association contain only partial signatures of species interactions, our study highlights that biotic homogenisation translates to finer changes in community structure by affecting the number, strength and type of species associations.

## 1. Introduction

Among the major effects of global change on biological diversity, the modification or even the extinction of species interactions has early on been identified as being pervasive, but is still poorly understood (Janzen, 1971; Diamond, 1989). Because there are many more interactions than species, a perturbation in species interactions may be decoupled from changes in species richness or community composition (Poisot *et al*., 2015; Gravel *et al*., 2019). In particular, modifications affecting species interactions can be stronger (Valiente-Banuet *et al*., 2015) or weaker (Li *et al*., 2018) than those affecting species richness. The structure and dynamics of species interactions are among the main drivers of community dynamics (Davis *et al*., 1998; Barabás *et al*., 2016), and therefore represent a critical subject of study for ecology and biodiversity conservation (Kissling & Schleuning, 2015; García-Girón *et al*., 2020). Despite the importance of integrating species interactions into conservation biology, we still have a limited understanding of the drivers and consequences of changes in the strength and the structure of species interactions.

In the last decades, there has been an increasing use of network approaches to study species interactions in empirical and theoretical communities (Bascompte *et al*., 2003; Ings *et al*., 2009; Kéfi *et al*., 2015; Trøjelsgaard & Olesen, 2016). Ecological communities can thus be depicted as interaction networks by defining nodes as individuals or species, and links between the nodes as species interactions (Newman *et al*., 2006). The estimation of species interactions is however subject to a conceptual question (*how are the strength and type of interactions defined?*) and a technical challenge (*how to estimate an interaction?*). In some cases (*e.g*. in simple trophic networks with few taxa), observations or experiments can address both issues as the existence and type of species interactions are clearly identified. However, these cases provide inference of species interactions in local and specific systems, compromising the possibility to derive general rules for interactions in larger communities (Whittaker *et al*., 2005; Denny & Benedetti-Cecchi, 2012). The empirical identification and measure of interactions in species-rich communities in particular, is challenged by the high number of potential interactions to be estimated (proportional to the square of species number) (Barner *et al*., 2018). An alternative approach is to assume that species *associations* (inferred from their spatial aggregation) are partly shaped, at least to some extent, by the combination of true interactions (*i.e*. clear ecological relationships such as competition or predation). In this case, studying communities with a large number of species and broad spatial coverage should be a good framework for estimating species associations, although the ability of spatial co-occurrence patterns to infer pairwise species interactions is still controversial (Blanchet *et al*., 2020).

Nonetheless, species co-occurrence might be an information-rich proxy of the outcome of direct and indirect biotic interactions in communities (Delalandre & Montesinos-Navarro, 2018; Freilich *et al*., 2018). Indeed, the composition of a local community results from interspecific interactions as well as multiple intertwined processes generating patterns of spatial aggregation between species (Fig. 1) (Wisz *et al*., 2013). These factors include neutral processes (regional dispersion and local stochasticity (Hubbell, 2001)), historical processes (phylogeography (Kraft *et al*., 2007)) and niche processes (HilleRisLambers *et al*., 2012; Letten *et al*., 2017). Niche processes combine what are sometimes referred to as Grinnellian and Eltonian processes (Chase & Leibold, 2003; Devictor *et al*., 2010a). Grinnellian processes (Grinnell (1917); later extended by Hutchinson, 1957) consider the niche as the species’ response to environmental conditions acting as an environmental filter for the community. Eltonian processes (Elton, 1927) consider the niche as the species’ impact on its environment and refers to the mutual dependency of species with each other, including the limiting similarity hypothesis (*i.e*. the niche overlap between two species that limits their coexistence) (MacArthur & Levins, 1967; Abrams, 1975; Martin & Bonier, 2018) and facilitation between species (*e.g*. cooperation, exchange of social information (Seppänen *et al*., 2007; Gil *et al*., 2019; Tu *et al*., 2019)).

**Figure 1:**
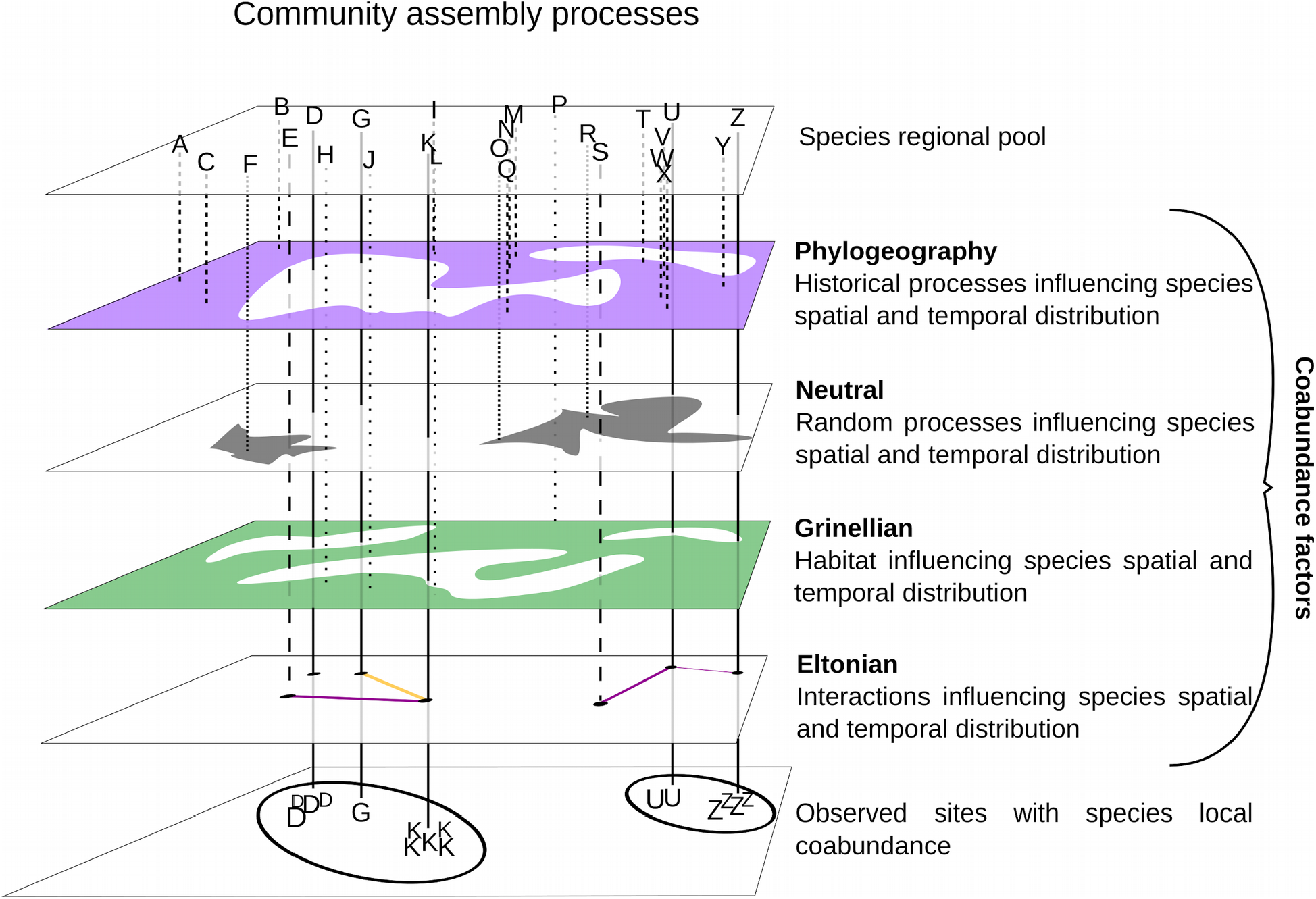
Community assembly processes and species co-abundance. Species interactions that influence species spatial aggregation (or segregation) and temporal change in abundance are referred to as the Eltonian component of species co-abundance. In addition to the Eltonian component, coabundances are also the result of habitat filtering (Grinellian), random processes due to neutral dispersal, as well as historical processes related to the species phylogeography. The result of all these processes leads to the observed species co-abundances. Each letter stands for a different species. Species U and Z share a common biogeographic region and random processes have not prevented them from co-occurring. As they live in a similar habitat and interact in a way that enables their coexistence, they can be observed together in the same location at the same time.

Refining the Eltonian component (*i.e*. the part of the co-occurrences due to the biotic filter) can be done through multiple approaches (Kissling *et al*., 2012). Based on null models, one can control for the species associations that are simply expected by chance rather than grounded in ecological processes by testing whether species are found together more or less frequently than expected by chance (Gotelli, 2000; Ulrich & Gotelli, 2010; Kohli *et al*., 2018). Indirect effects between species (*i.e*. the effect of a third species on the association between two other species) can be evaluated using partial correlations (Faust & Raes, 2012; Harris, 2016). Recent progress with Joint Species Distribution Models have also provided ecologists with new tools for estimating species associations by studying residual co-occurrence patterns after accounting for environmental niches from large data-sets (Tikhonov *et al*., 2017; Zurell *et al*., 2018). Overall, recent methods removing non-Eltonian components from co-occurrences (Azaele *et al*., 2010; Faisal *et al*., 2010; Ovaskainen *et al*., 2010; Lindenmayer *et al*., 2015) are promising for uncovering species association networks (Araújo *et al*., 2011; Morueta-Holme *et al*., 2016). Co-occurrences are thus information-rich proxies of the outcome of direct and indirect biotic interactions in communities (Delalandre & Montesinos-Navarro, 2018; Freilich *et al*., 2018). In particular, if interaction networks remain out of reach (Sander *et al*., 2017; Thurman *et al*., 2019), association networks based on species co-occurrence may be useful for capturing community organisation through aggregated community indices, *i.e*. statistics summarising an aspect of the network at the community level (Barner *et al*., 2018).

Tracking large-scale changes in species associations might represent a significant advance for macro-ecology and conservation biogeography. Indeed, ecological processes are ultimately influenced by which and how species interact (Cardinale *et al*., 2002; Goudard & Loreau, 2008). Moreover, the responses of species associations to environmental changes are not necessarily proportional to the responses of individual species (Valiente-Banuet *et al*., 2015). Therefore, measuring community changes through the change in species diversity within local communities or, at a larger scale, between communities (for instance using β-diversity) may mask important modifications of the structure and properties of those communities (Poisot *et al*., 2017). For instance, the replacement of a set of diverse and mainly specialist species by a few generalists (McKinney & Lockwood, 1999; Olden *et al*., 2004) is a well-documented form of biotic homogenisation (Clavel *et al*., 2011). In those communities, the composition tends to be closer to random expectations (Barnagaud *et al*., 2017), *i.e*. with less and less visible niches processes. Yet, anthropogenic perturbations can also act as a strong filter selecting for more specialist species (Gaüzère *et al*., 2020), in which case the Grinellian filter outweighs the Eltonian one. While many studies have evidenced the impact of global change on biotic homogenisation (Newbold *et al*., 2018), whether homogenisation in species composition is related to a directional change in species associations remains to be explored (Li *et al*., 2018).

In this study, we conducted a large-scale spatio-temporal analysis of bird species association networks. Birds form a relevant group to estimate ecologically meaningful species associations. Indeed, the Eltonian filter is likely to impact bird co-occurrences, as among other interacting processes, competitive exclusion as well as social information exchanges have been frequently shown to occur between birds (Thomson *et al*., 2003; Forsman & Thomson, 2008; Magrath *et al*., 2015). Furthermore, the fact that bird species have been widely monitored for decades allows to track changes in species association patterns in space and time; an opportunity that is not possible for other groups of species. Using data from the French Breeding Bird Survey, we addressed the following two objectives:

1. *reconstruct species association networks in communities from co-abundance data*. To do so, we first inferred species associations from co-abundance (co-occurrence with abundance) data (Fig. 2a) corrected for non-Eltonian co-occurrence processes. We then quantified different aspects of the species association networks using three complementary network indices: intensity, attractiveness and clique structure of the network. Intensity corresponds to the mean association strength. Attractiveness is the ratio of positive/negative associations, and clique structure describes the structural complexity of the association network (Fig. 2b).
2. *test whether biotic homogenisation was linked to directional changes in association networks*. We analysed the relationships between the spatio-temporal dynamics in bird network indices and the spatio-temporal dynamics in β-diversity, at a large scale and separately in three main types of habitats, woodland, grassland and human settlements. We also tested whether habitat generalist and specialist species were responsible for the observed changes in species associations over time and space.

**Figure 2:**
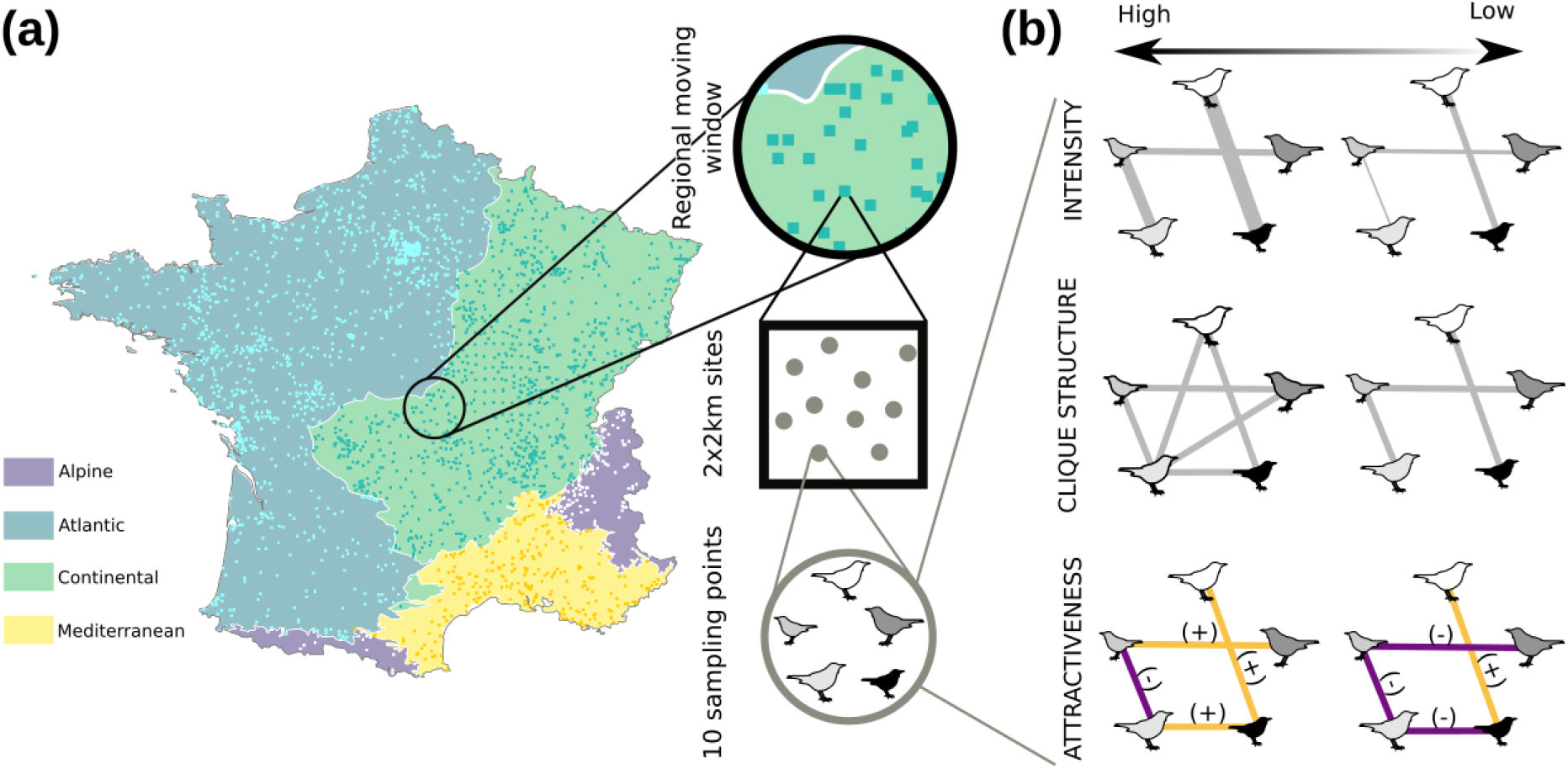
From bird monitoring to association network indices. (a) Spatial distribution of the 1,969 selected (out of 2,514) sites from the 2001 to 2017 FBBS (STOC-EPS). On each site (2×2 km square), bird observations were recorded on 10 sampling points. Geographically Weighted Regression using data from moving windows were used to assess metric values. Mainland France was split into 4 biogeographic regions (Alpine, Atlantic, Continental and Mediterranean). (b) Association network indices: intensity (*i.e*. mean strength of species associations), attractiveness (predominance of positive (light orange) or negative (dark purple) species associations) and clique structure (level of structuration of the species association network). Lines represent species associations. For each index, examples corresponding to high and low values are displayed from left to right, respectively. For intensity, the thicker the line, the higher the absolute value of the association. For attractiveness, the high value example is 0.5 (three positive associations and one negative association out of 4 existing associations) and the low value is −0.5. For clique structure, the high value is 0.125 (two realised cliques out of 16 possible cliques) and the low value is 0.

## 2. Materials and methods

### 2.1 Bird data

Bird data were extracted from the French Breeding Bird Survey (FBBS) (Jiguet *et al*., 2012). In this scheme, volunteer ornithologists monitored common bird species on 2,514 sites (Fig. 2a) from 2001 to 2017, following a standardised protocol. Sites are 2×2 km squares in which abundances of breeding bird species were monitored on 10 homogeneously distributed sampling points across habitats in the landscape. In order to avoid habitat classes with too few observations, we grouped the 37 main types of habitat described in the field in 19 classes for the estimation of species associations, and in three groups, corresponding to the three main types of habitats of our dataset (woodland, grassland and human settlements), for network analyses. Among the 242 species recorded in the dataset, we selected the 109 most abundant species (representing 99% of the total abundance) to avoid any over-representation of rare species (that are therefore more difficult to monitor). After removing rare species and the sites only monitored once, our dataset comprised 19,580 sampling points in 1,969 sites and 109 species (for habitat and species details, see Appendix 1 in Supporting Information).

#### 2.2 Species associations

We estimated associations between pairs of species from bird co-abundance data (Morueta-Holme *et al*., 2016) for each year (2001 to 2017), for each of the four biogeographic regions and for each of the 19 habitats using the five following steps (Fig. 3).

**Figure 3:**
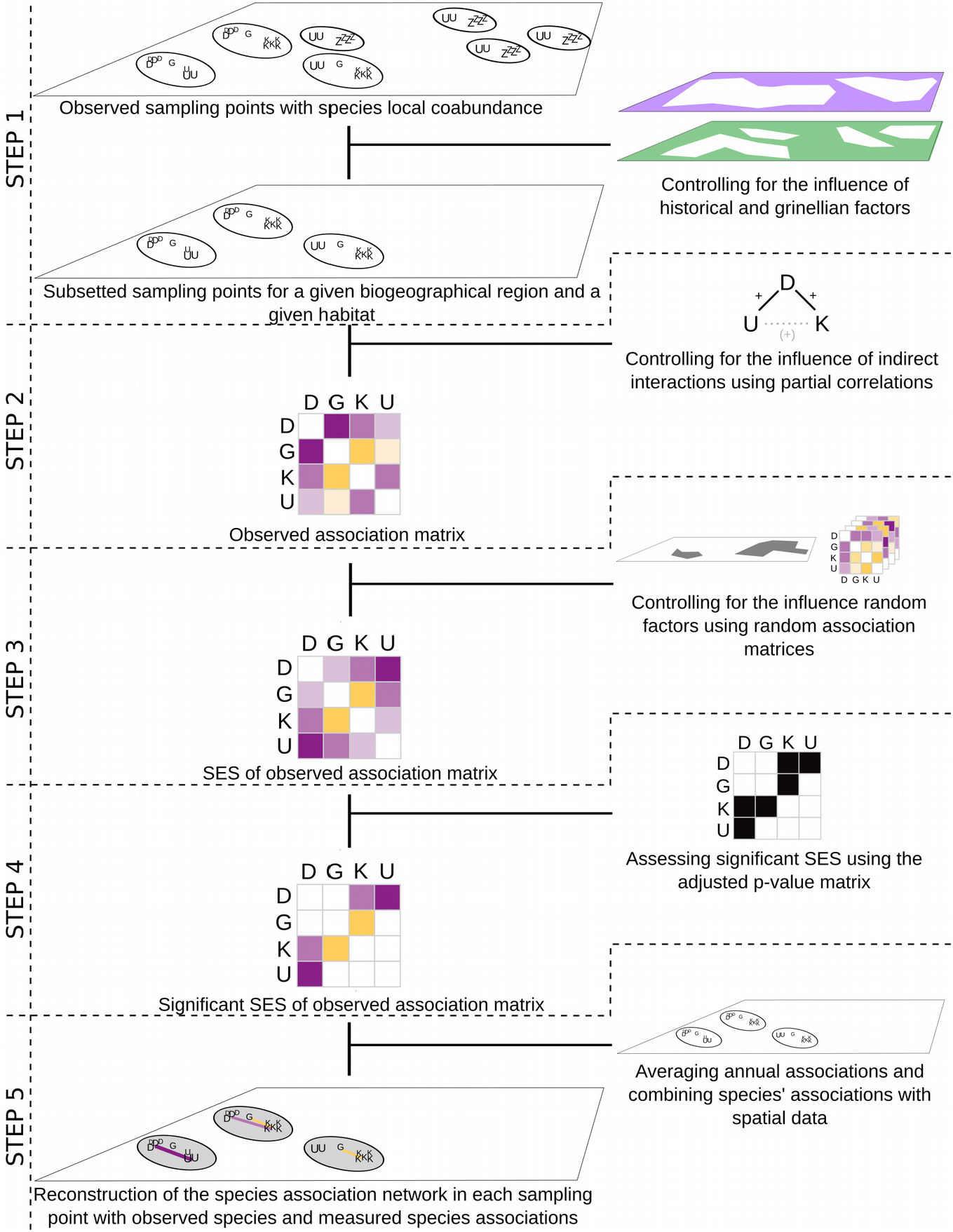
Workflow for estimating species association networks. For each biogeographical region and each habitat (Step 1), an observed association matrix was obtained by partial correlations (Step 2) (positive correlations in orange, negative correlations in purple, colour intensity proportional to correlation strength) from co-abundance data (species D, K, G, U as an example). Random association matrices were calculated using partial correlations on permuted datasets (1000 times) and were used to calculate standard effect sizes (Step 3) of observed associations as well as their adjusted p-values to obtain significant SES of the observed association matrix (Step 4). Steps 1 to 4 were repeated for each year providing annual associations, which were then averaged over years for each species pair. species associations were finally added to the spatial co-abundance data to obtain a species association network for each of the sampling points (Step 5).

Step 1. In order to limit the influence of phylogeography and habitat features on species associations, we first grouped the data by biogeographic region (Continental, Atlantic, Mediterranean, Alpine), by habitat and by year to estimate an association for each pair of bird species, for each year, for each of the four biogeographic regions (EEA, 2016) and for each of the 19 habitat classes. Note that we used the most detailed habitat information available but, as finer habitat grain is out of reach, we admit that not all the influence of the Grinnellian filter has been removed by this first grouping.

Step 2. In each biogeographic region and habitat, we used the log-transformed co-abundance data (to obtain normally distributed data) to calculate observed associations as partial correlations between each pair of species (Schäfer & Strimmer, 2005) as follows (Eq. 1):

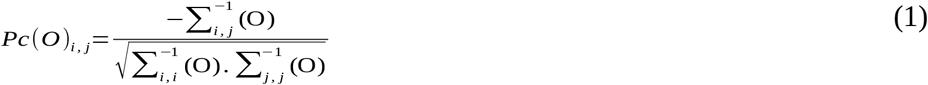

with *O* the matrix of observed abundance (species x sites), *Pc*(*O*)_*i,j*_ the partial correlation between species *i* and *j*, and 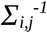 the value for species *i* and *j* of the inverse of the covariance matrix. Given that the association between two species can be influenced by the presence of another co-occurring species, this approach partially removes the indirect effects of the other co-occurring species on the estimated association between the two considered species by focusing on the conditional association (Harris, 2016).

Step 3. Partial correlations can be affected by species commonness, since common species have higher probabilities to co-occur than less abundant species because of a higher representativeness in the data (Blüthgen *et al*., 2008). To correct this bias, we computed partial correlations on 1000 random co-abundance datasets obtained by keeping constant the total number of individuals in a given sampling point, and assuming that the probability for a species to occur in a given sampling point was proportional to its frequency in the dataset. We then calculated partial correlation standardised effect size of species *i* and *j* (*SES_i,j_*) as follows (Eq. 2):

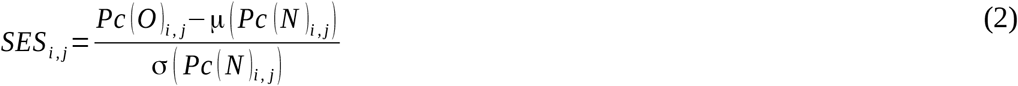

where *Pc*(*O*)_*i,j*_ is the observed partial correlation between species *i* and species *j, μ*(*Pc*(*N*))_*i,j*_ and *σ*(*Pc*(*N*))_*i,j*_ the mean and standard deviation of partial correlations from the 1000 randomly sampled datasets.

Step 4. In order to identify *significant* associations, we calculated a two tail p-value for each pairwise association using the rank of the observed association in the Gaussian distribution of null associations obtained from step 3. That is, we determined the number of replicates for which the absolute value of the observed partial correlation is greater than the absolute null partial correlation (p-values were corrected for multiple comparisons following Benjamini & Hochberg (1995)). Significant associations therefore corresponded to *SES_i,j_* for which adjusted p-values were below 0.05.

Step 5. For each species pair, for each biogeographic region and for each habitat, we averaged the significant associations over the 17 annual associations (one for each year). In the absence of any significant association across the 17 years, the association was considered null (*i.e*. equal to zero). This results in a total set of 260,191 association estimates, spread over 5,886 pairs of species, four biogeographic regions and 19 habitats.

#### 2.3 Association network indices

We considered three mathematically independent indices that describe different aspects of the network built upon pairwise association estimates (Fig. 2b and see examples in Appendix 2). The three indices were calculated for 121,172 networks corresponding to the communities monitored in the 19,580 sampling points between 2001 and 2017.

Intensity *I* quantifies the strength of associations in the species association network of a community. It reflects the average intensity of the associations in the network. It is weighted to account for the differences between species’ abundances (Eq. 3).

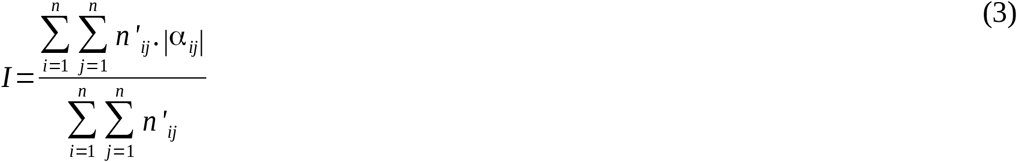

with *n* the number of species, *n*’_*ij*_ the number of pairs of species *i* and *j* in the community pool, and *α*_*ij*_ the association (as defined in step 5) between species *i* and *j* (with *i*≠*j*; when *i*=*j, α*_*ij*_=0). *I* varies between 0 and |*α*|_max_. High values of *I* are reached in communities including mainly strong associations.

Attractiveness *A* quantifies the prevalent sign of the associations as the number of positive associations minus the number of negative associations standardised by the total number of associations (Eq. 4). Attractiveness is analogous to the *association ratio* in plant networks (Saiz *et al*., 2014). However we choose to label this metric *attractiveness* rather than *association ratio* because it also stands for methods estimating associations (Chiyo *et al*., 2011).

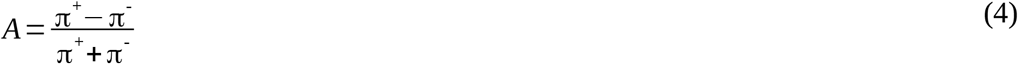

with π^+^ the number of positive associations and π^-^ the number of negative associations. It varies between −1 (if all the associations are negative) and 1 (if all the associations are positive).

Clique structure *C* quantifies the level of structuring of the species association network. It is calculated using the number of existing cliques (*i.e*. fully connected groups of species, (Luce & Perry, 1949)) with three or more nodes, standardised by the number of potential cliques in a given network (Eq. 5).

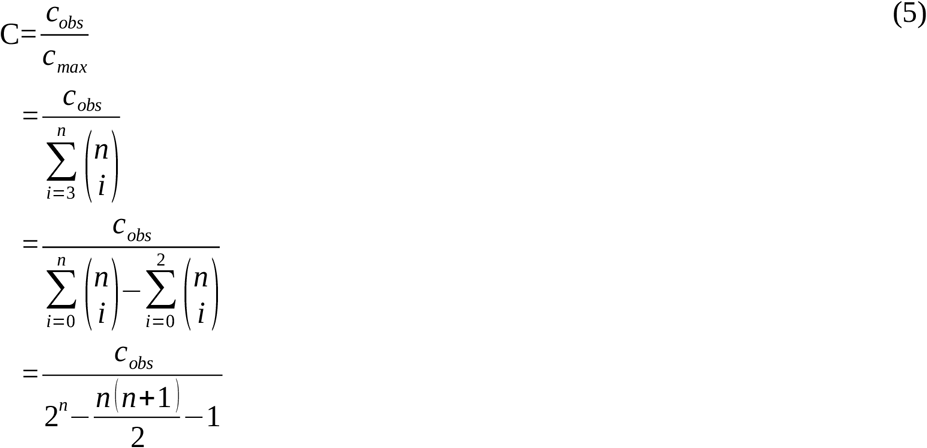

with *c_max_* the maximum possible number of 3- to *n*-cliques, *c_obs_* the observed number of 3- to *n*-cliques, *n* the number of species in the network.

*C* quantifies the complexity of the network architecture resulting from the interweaving of associated species (see Appendix 2). Networks with high *C* values have a complex structure, with multiple imbricated groups of interconnected species. Networks with low *C* only have a few small sized interconnected groups of species.

#### 2.4 Spatial averages and temporal trends in association network indices

##### 2.4.1 Spatial averages of association network indices

For spatial analyses, we averaged the annual values of each index (*I, A* and *C*) for each sampling point, resulting in one value for each index for each sampling point. We then computed the *spatial window values* of each association network index, for each site and for each year, using an 80-km radius window. We determined the window size as a compromise between a large spatial coverage and a fine spatial resolution and we conducted all subsequent analyses for various radii to assess the robustness of our results to changes in the window size (see Appendix 3). Spatial window values were computed to analyse, on a similar spatial scale, the relationships between community indices and β-diversity which is an inter-site measure based on species data from multiple sites (see below part 2.5). It also provided more complete data when sampling points or sites were not monitored every year, in particular for calculating temporal trends (see below part 2.4.2). We estimated spatial window values using Geographically Weighted Regression (GWR) (Brunsdon *et al*., 1996; Gollini *et al*., 2015). In this approach, the centre of each site was consecutively considered as the centre of a fixed radius window. Each index was calculated using data from all sampling points encapsulated within the spatial window. A weight was attributed to each sampling point, which decreased with the distance to the central selected site following a bisquare kernel function.

##### 2.4.2 Temporal trends of association network indices

We estimated the *spatial window trends* as the temporal trend of each association network index (*I, A* and *C*) following the same framework as for spatial window values (see above 2.4.1). The trend of each index corresponds to the coefficient of a linear regression calculated using annual index values in the selected sampling points, weighted according to their proximity to the central site. Only significant trends were used for subsequent analyses.

In addition to being calculated on the whole dataset, spatial and temporal window values were also computed for each of the three mains groups of habitat (woodland, grassland and human settlements). In this case, we selected sampling points that 1) belonged to the type of habitat considered, and 2) were located within sites dominated by this type of habitat. We recalculated spatial averages of network indices (too few trends were significant to conduct temporal analyses) using the subset of sampling points for each of the three main groups of habitat.

#### 2.5 β-diversity and Community Generalisation Index

##### 2.5.1 Spatial and temporal variation

We assessed the spatial and temporal variations of β-diversity and proportion of habitat generalists following the same framework as for spatial window values (see paragraph 2.4.1). β-diversity corresponds to species diversity between a set of sites and is the result of two components, *species turnover sensu stricto* and *nestedness* (Baselga, 2010). A raw observed species turnover between two communities, A and B, can results from a simple difference in community size if B includes fewer species. In this case, part of the observed turnover is due to the nested composition of B in the composition of A. That is, a simple species loss from A to B generates a turnover due to a variation in community size. In contrast, the species turnover *s.s*. corresponds to the species not shared by the two communities, *i.e*. resulting from the replacement of one species by another. We first aggregated species data from sampling point level to site level to obtain species data for each site. We then randomly selected 10 sites in each spatial window (see 2.4.1) (Devictor *et al*., 2010b) and computed the species turnover *s.s*. (hereafter referred to as *β-diversity*) of the set of sites in that window using the *betapart* R package (Baselga & Orme, 2012). We repeated this selection step 10 times, and we took the mean of β-diversity.

We used the Species Generalisation Index (SGI) from Godet et al. (2015) to calculate for each community the Community Generalisation Index (CGI). SGI corresponds to the habitat specialisation value of a species independently from its abundance and range, and CGI to the average habitat specialisation of species in a given assemblage, weighted by their local abundances.

##### 2.5.2 Association network indices vs. β-diversity and CGI

We analysed the spatial relationships between β-diversity and the three association network indices as well as the spatial relationships between the CGI and network indices by performing general additive models (GAM) to assess the linear relationship between each of the three network indices and β-diversity or CGI, while explicitly modelling the spatial autocorrelation. That is, to test the link between spatial values of community indices and β-diversity, each association network index (*I, A* and *C*) was successively considered as the response variable regressed over β-diversity or CGI. We explicitly modelled the spatial autocorrelation using a two dimension isometric thin plate regression spline based on geographic coordinates of sites following Wood (2003, 2017). As species richness may influence network indices, we added species richness as an explanatory variable to disentangle the effect of β-diversity or CGI from the effect of species richness on network indices. In addition, as part of the results could be driven, at least to some extent, by unchecked structural relationships between association indices and beta diversity, we therefore tested whether the relationships obtained could be due to the intrinsic redundancy between network indices and β-diversity using simulated and permuted data (see Appendix 4).

Using a similar model, we also tested the temporal relationship between the trends in the three association network indices and the trend in β-diversity or CGI (accounting for the trend in species richness) using their spatial window trends. Limits of relying on space-for-time substitution (*i.e*. relying only on spatial gradient to infer temporal relationships) are well documented (Damgaard, 2019), this final step was therefore essential to support results from the spatial analysis. Due to data limitation, this temporal analysis was only possible at the national scale and not in each of main type of habitat.

## 3. Results

### 3.1 Species associations from co-abundance

We found 8.1% of positive associations, 38.3% of negative associations, whereas 53.6% of associations were non-significant. 40% of the species pairs showed qualitatively constant associations (*i.e*. significant associations that were positive or negative in more than 90% of cases) across habitat/biogeographic region combinations. On average, each species was associated with 1 to 93 other species (mean=41, sd=26) with variations between habitats and biogeographic regions (associations available in Appendix 5). In particular, in woodland, we found 4.3% of positive associations (*e.g*. between the Eurasian wryneck (*Jynx torquilla*) and the lesser spotted woodpecker (*Dryobates minor*)), 28.7% of negative associations (*e.g*. between the common chaffinch (*Fringilla coelebs*) and the common cuckoo (*Cuculus canorus*)) and 67.0% of non-significant associations. In grassland, we found 7.8% of positive associations (*e.g*. between the calandra lark (*Melanocorypha calandra*) and the Eurasian skylark (*Alauda arvensis*)), 29.1% of negative associations (*e.g*. between the red-legged partridge (*Alectoris rufa*) and the grey partridge (*Perdix perdix*)) and 63.1% of non-significant associations. In human settlements, we found 4.6% of positive associations (*e.g*. between the house sparrow (*Passer domesticus*) and the Eurasian collared dove (*Streptopelia decaocto*)), 30.1% of negative associations (e.g. between the house sparrow (*Passer domesticus*) and the carrion crow (*Corvus corone*)) and 64.7% of non-significant associations.

### 3.2 Overall relationship between association network indices, β-diversity and CGI in space and time

In space, intensity was positively related to β-diversity (Fig. 4a) and negatively to CGI (Fig. 4d). Attractiveness was negatively related to β-diversity (Fig. 4b) and not significantly related to CGI (Fig. 4e). Clique structure was positively related to β-diversity (Fig. 4c) and negatively to CGI (Fig. 4f). Model outcomes are reported in Table 1.

**Figure 4:**
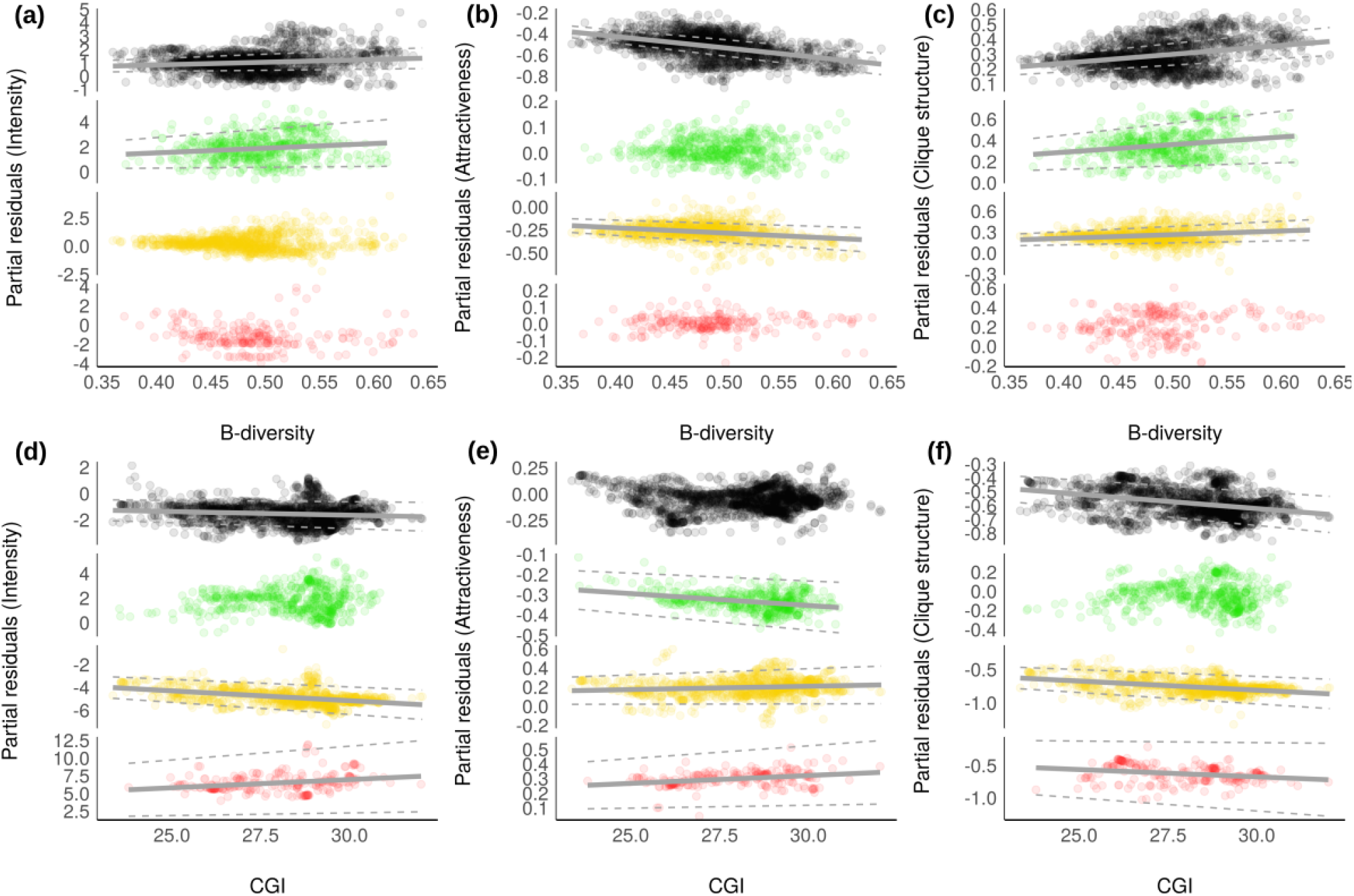
Relationships between network indices versus β-diversity and CGI for all habitats (black dots), woodland (green dots), grassland (yellow dots) and human settlements (red dots). First row: relationships between (a) intensity and β-diversity, (b) attractiveness and β-diversity, (c) clique structure and β-diversity. Second row: relationship between (d) intensity and CGI, (e) attractiveness and CGI, (f) clique structure and CGI. Dots correspond to partial residuals of the regression models regressed over predictors and regression lines (solid lines) with confidence intervals (dashed lines) are shown when significant.

**Table 1:**
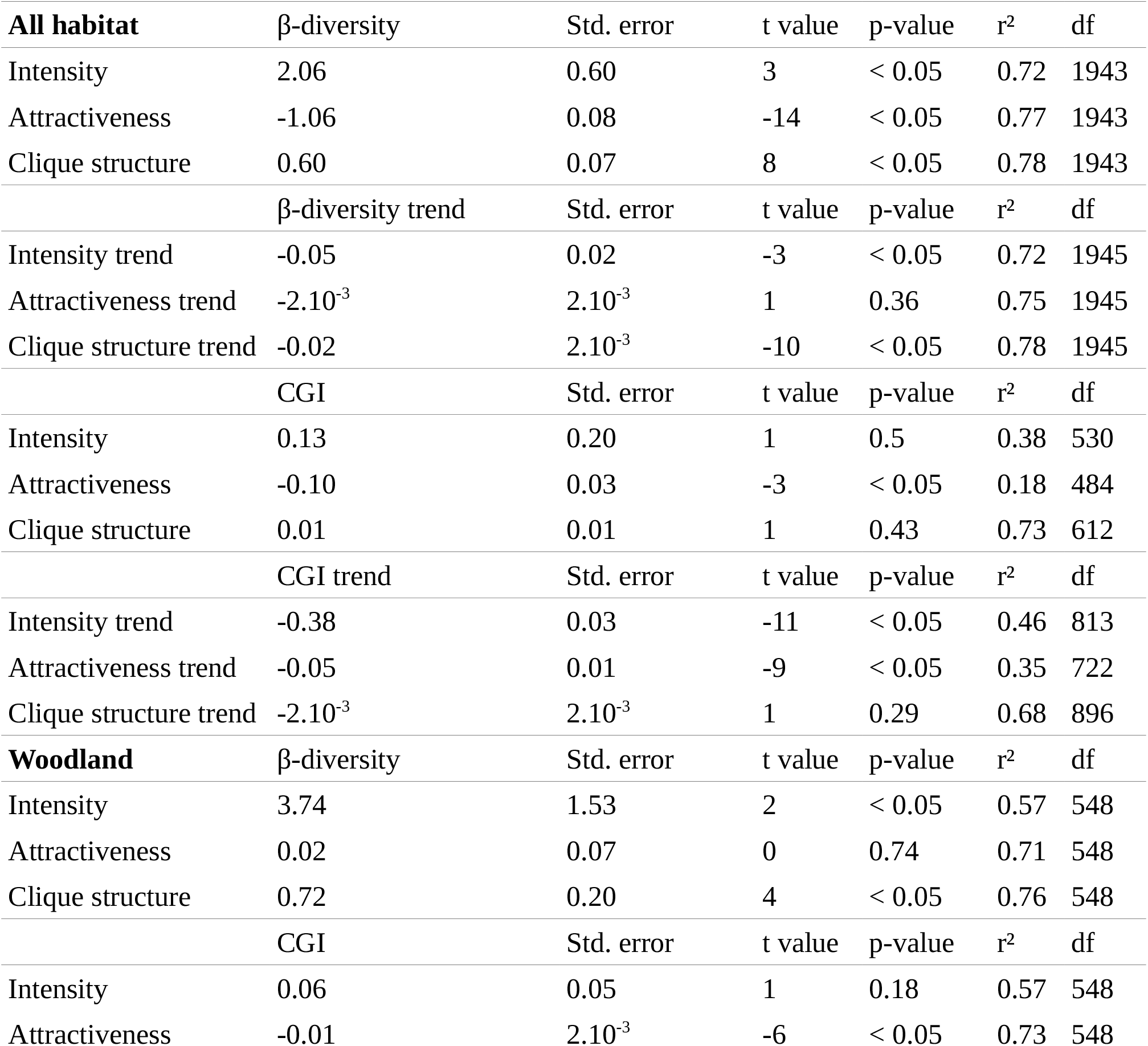
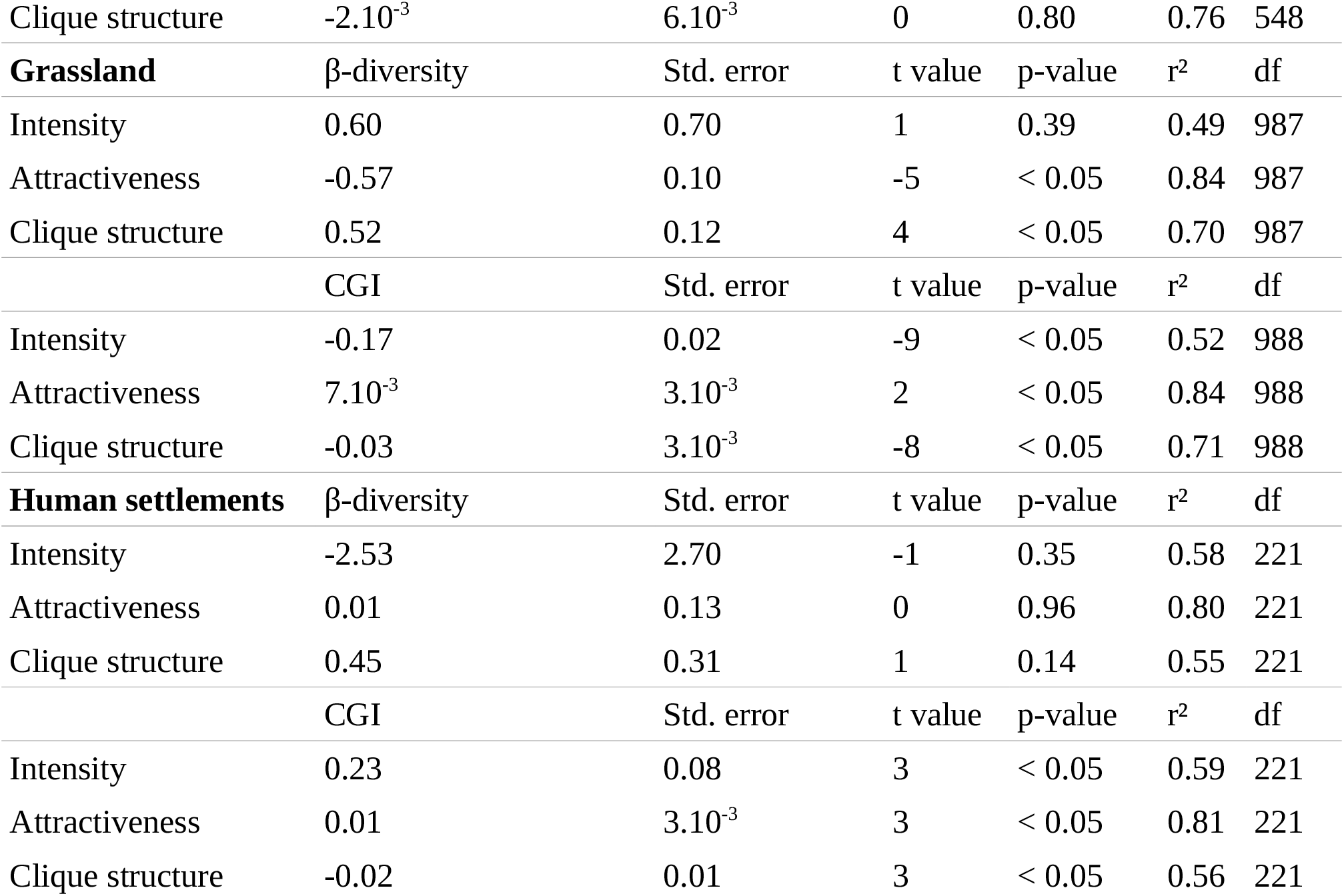
Result summary of the GAM models, coefficient estimates (*β-diversity* or *CGI*), standard errors (*Std. error*), associated *t value*, significance level (*p-value*), goodness of fit (*r*^2^) and degree of freedom (*df*).

| **All habitat** | β-diversity | Std. error | t value | p-value | r<sup>2</sup> | df |
| --- | --- | --- | --- | --- | --- | --- |
| Intensity | 2.06 | 0.60 | 3 | < 0.05 | 0.72 | 1943 |
| Attractiveness | -1.06 | 0.08 | -14 | < 0.05 | 0.77 | 1943 |
| Clique structure | 0.60 | 0.07 | 8 | < 0.05 | 0.78 | 1943 |
| | β-diversity trend | Std. error | t value | p-value | r<sup>2</sup> | df |
| Intensity trend | -0.05 | 0.02 | -3 | < 0.05 | 0.72 | 1945 |
| Attractiveness trend | -2.10<sup>-3</sup> | 2.10<sup>-3</sup> | 1 | 0.36 | 0.75 | 1945 |
| Clique structure trend | -0.02 | 2.10<sup>-3</sup> | -10 | < 0.05 | 0.78 | 1945 |
|  | CGI | Std. error | t value | p-value | r<sup>2</sup> | df |
| Intensity | 0.13 | 0.20 | 1 | 0.5 | 0.38 | 530 |
| Attractiveness | -0.10 | 0.03 | -3 | < 0.05 | 0.18 | 484 |
| Clique structure | 0.01 | 0.01 | 1 | 0.43 | 0.73 | 612 |
|  | CGI trend | Std. error | t value | p-value | r<sup>2</sup> | df |
| Intensity trend | -0.38 | 0.03 | -11 | < 0.05 | 0.46 | 813 |
| Attractiveness trend | -0.05 | 0.01 | -9 | < 0.05 | 0.35 | 722 |
| Clique structure trend | -2.10<sup>-3</sup> | 2.10<sup>-3</sup> | 1 | 0.29 | 0.68 | 896 |
| **Woodland** | β-diversity | Std. error | t value | p-value | r<sup>2</sup> | df |
| Intensity | 3.74 | 1.53 | 2 | < 0.05 | 0.57 | 548 |
| Attractiveness | 0.02 | 0.07 | 0 | 0.74 | 0.71 | 548 |
| Clique structure | 0.72 | 0.20 | 4 | < 0.05 | 0.76 | 548 |
|  | CGI | Std. error | t value | p-value | r<sup>2</sup> | df |
| Intensity | 0.06 | 0.05 | 1 | 0.18 | 0.57 | 548 |
| Attractiveness | -0.01 | 2.10<sup>-3</sup> | -6 | < 0.05 | 0.73 | 548 |
| Clique structure | -2.10<sup>-3</sup> | 6.10<sup>-3</sup> | 0 | 0.80 | 0.76 | 548 |
| **Grassland** | β-diversity | Std. error | t value | p-value | r<sup>2</sup> | df |
| Intensity | 0.60 | 0.70 | 1 | 0.39 | 0.49 | 987 |
| Attractiveness | -0.57 | 0.10 | -5 | < 0.05 | 0.84 | 987 |
| Clique structure | 0.52 | 0.12 | 4 | < 0.05 | 0.70 | 987 |
|  | CGI | Std. error | t value | p-value | r<sup>2</sup> | df |
| Intensity | -0.17 | 0.02 | -9 | < 0.05 | 0.52 | 988 |
| Attractiveness | 7.10<sup>-3</sup> | 3.10<sup>-3</sup> | 2 | < 0.05 | 0.84 | 988 |
| Clique structure | -0.03 | 3.10<sup>-3</sup> | -8 | < 0.05 | 0.71 | 988 |
| **Human settlements** | β-diversity | Std. error | t value | p-value | r<sup>2</sup> | df |
| Intensity | -2.53 | 2.70 | -1 | 0.35 | 0.58 | 221 |
| Attractiveness | 0.01 | 0.13 | 0 | 0.96 | 0.80 | 221 |
| Clique structure | 0.45 | 0.31 | 1 | 0.14 | 0.55 | 221 |
|  | CGI | Std. error | t value | p-value | r<sup>2</sup> | df |
| Intensity | 0.23 | 0.08 | 3 | < 0.05 | 0.59 | 221 |
| Attractiveness | 0.01 | 3.10<sup>-3</sup> | 3 | < 0.05 | 0.81 | 221 |
| Clique structure | -0.02 | 0.01 | 3 | < 0.05 | 0.56 | 221 |

In time, the temporal trend in intensity was not significantly related to the temporal trend in β-diversity but negatively related to the trend in CGI. The temporal trend in attractiveness was negatively related to the trend in β-diversity and to the temporal trend in CGI. The trend in clique structure was not significantly related to the trend in β-diversity nor to the trend in CGI. These temporal results corroborated the relationships found above with spatial values for attractiveness but not for intensity or clique structure.

### 3.3 Relationship between association network indices, β-diversity and CGI by habitat

In woodland, intensity was positively related to β-diversity but not to CGI (Tab. 1, Fig. 4). Attractiveness was not significantly related to β-diversity and negatively related to CGI. Clique structure was positively related to β-diversity but not related to CGI.

In grassland, intensity was not significantly related to β-diversity and negatively related to CGI. Attractiveness was negatively related to β-diversity and positively related to CGI. Clique structure was positively related to β-diversity and negatively related to CGI.

In human settlements, intensity was not significantly related to β-diversity and positively related to CGI. Attractiveness was not significantly related to β-diversity but positively related to CGI. Clique structure was not significantly related to β-diversity but negatively related to CGI.

## 4. Discussion

Biotic homogenisation (*i.e*. the replacement of a diversity of mainly specialist species by a few generalists (McKinney & Lockwood, 1999)) triggered by ongoing global change (Lockwood *et al*., 2000; Devictor *et al*., 2008; Godet *et al*., 2015) is considered as one of the most pervasive aspects of the biodiversity crisis (Olden *et al*., 2004). At the local scale, we measured the homogenisation of bird communities as a decrease of β-diversity (McGill *et al*., 2015) in space and time while considering changes in the proportion of habitat generalists. Our study unravelled clear relationships between species homogenisation and changes affecting species associations. These relationships could be revealed thanks to the reconstruction of association networks from co-abundance data and to the ability of tracking modifications in the structure of those association networks. We showed that biotic homogenisation was linked to weaker intensity and clique structure, and with more positive attractiveness. In other words, more similar areas in terms of species composition sheltered weaker and relatively more positive associations but less structured association networks.

Several studies have recurrently shown the difficulties to use species associations as reliable proxies for species interactions (Sander *et al*., 2017; Freilich *et al*., 2018; Blanchet *et al*., 2020). Species associations are indeed potentially affected by non-biotic filters and some types of species interactions remain inaccessible from co-occurrence, *e.g*. amensalism (Morales-Castilla *et al*., 2015). While our methodology takes into account non-biotic filters, it is still subject to remnant effects of those filters and additional processes linked to life-history traits (*e.g*. dispersal abilities). That means that some of the species associations we found are still likely to result from, for instance, fine grain habitat filtering, *e.g*. the negative association between the short-toed treecreeper and the Eurasian robin is probably due to the preference of the latter for young forest whereas the former is rather found in old stands (Laiolo *et al*., 2004). Another pitfall is the difficulty to estimate temporal variation in species associations from co-occurrence data as, currently, only state-space models may allow to quantify species interrelations in varying environments (Deyle *et al*., 2016) and this approach requires long time-series generally not available across multiple sites and at large scales. This prevented us from estimating temporal variations in associations, although species interactions are known to vary in the short (Price *et al*., 2005; Olesen *et al*., 2008) and long term (Li & Waller, 2016; Lyons *et al*., 2016) particularly in response to environmental changes (Rico-Gray *et al*., 2012; Bimler *et al*., 2018; Clark *et al*., 2018).

In spite of these limitations, our approach was able to capture pairwise associations that could be related to existing knowledge on bird behaviour and interactions. For instance, the negative associations inside the *Parus* guild, in particular between the goldcrest (*Regulus regulus*) and titmice (*Poecile montanus, Lophophanes cristatus*) may be related to dominance behaviour of the last two species leading to the spatial exclusion of the goldcrest (Alatalo *et al*., 1985). The negative association between the common chaffinch and the common cuckoo can be linked to the chaffinch’s choice for breeding-site location that reduces parasitism from the cuckoo (Tolvanen *et al*., 2017). Negative associations between the mistle thrush (*Turdus viscivorus*) and the common blackbird or the great spotted woodpecker (*Dendrocopos major*) can be the result of territoriality as reported by Skórka & Wójcik (2005), whereas the negative association between the red-legged and the grey partridge are in line with the interspecific competition reported by (Rinaud *et al*., 2020). The positive association between the Eurasian treecreeper (*Certhia familiaris*) and the goldcrest may result from exploitation of the treecreeper using high vigilance information from *Parus* guild species (Henderson, 1989). The positive association between the Eurasian wryneck and the lesser spotted woodpecker can be related to the reuse of lesser spotted woodpecker’s cavities by the wryneck (Pakkala *et al*., 2019). The calandra lark and the Eurasian skylark, positively associated, are known to reciprocally attract each other (Delgado *et al*., 2013), as well as the house sparrow and the Eurasian collared dove (Skórka *et al*., 2016). In addition to these examples, association network indices that we used correspond to aggregated indices that are likely to provide a useful proxy to explore the drivers of changes in ecological communities, even considering the gap between species associations and species interactions (Barner *et al*., 2018). We are therefore convinced that species associations encapsulate relevant information about the structure of these communities and its changes in space and time.

Networks of detailed interactions between birds remained limited to local communities (Orchan *et al*., 2013) and association networks to woodland assemblages (Lane *et al*., 2014; Mokross *et al*., 2014). Our results including communities from different habitats at large scale emphasize that biotic homogenisation and modifications in association networks are not independent processes, which bring about a new repercussion of environmental change and species community homogenisation (see also Li et al. (2018)). Overall, intensity decreases with biotic homogenisation (intensity increases with β-diversity and declines with CGI) implying that homogenised communities are mainly composed of habitat generalist species weakly associated with each other. The link between β-diversity and CGI (see Appendix 6) corresponds to an overall pattern visible at the European level (Le Viol *et al*., 2012) in which species contributing to increase the community similarity are more likely to be habitat generalists. Moreover, as the attractiveness decreases with β-diversity, it implies that in the differentiated communities (as opposed to homogenised communities), negative associations are predominant. This could result, in particular, from competitive behaviour for instance for nest location or food as previously found in bird communities (Orchan *et al*., 2013; Lane *et al*., 2014). Finally, clique structure decreases with biotic homogenisation (clique structure increases with β-diversity and declines with CGI), suggesting that differentiated communities tend to have more complex network structures than homogenised communities. In other words, as the Eltonian filter is less and less visible, biotic homogenisation shapes sparse association networks in communities with weakly associated generalists.

However, patterns found between network indices and β-diversity were not similar among all habitats. In forest areas, β-diversity was higher than in other habitats and not related to the relative abundance of generalists versus specialists (Appendix 6). This indicates that forest communities were mostly composed of species with the same level of specialisation, as previously found in the French avifauna (Julliard *et al*., 2006). More particularly, specialists were together with other functionally close specialists and generalists with other functionally close generalists (see details in Appendix 7). In those differentiated (either specialised or generalised) forest communities, associations were stronger and formed more complex networks. However, associations were more negative in communities with habitat generalists than in communities with specialists implying that social information should be more important among habitat specialists and competition among habitat generalists.

Conversely, in more perturbed areas, such as human settlements, association intensity and β-diversity were not related. Instead, intensity was negatively related to the amount of habitat specialists. The most differentiated communities were shaped by the strong environmental filter formed by the urban environment. This selected mostly specialist species, considered as urban winners (Guetté *et al*., 2017), forming differentiated communities of species able to deal with this environment. Such a filter selecting for specialists in perturbed areas was previously shown on bird species (Gaüzère *et al*., 2020). But these specialists were functionally far from each other and, consequently, they did not strongly interact as they did not “know” the other species (Mönkkönen *et al*., 2017). Social information that can be shared by those species is therefore limited, explaining the low attractiveness observed. This leads to a scenario in which differentiated communities with specialists are composed of weakly and negatively associated species, selected for their ability to subsist in an environment strongly modified by humans, forming sparse association networks. In other words, the Grinellian filter appears to strongly outweigh the Eltonian filter in those communities. Grassland communities had intermediate levels of diversity and specialisation compared to the two other types of habitats. Intensity of associations was not linked to species diversity but to the amount of habitat specialists. Specialists, in differentiated communities, were also more negatively associated and formed more complex association networks. Biotic homogenisation in grassland seems therefore to be at the expense of the competing habitat specialists while reshaping association networks toward weakly and less negatively associated generalist species forming sparse association networks.

In short, exploring the fate of species associations provides a new dimension to the biotic homogenisation process, in addition to the homogenisation of species composition as homogenised communities have, in general, weaker, less negative and more simple association networks. Using species associations could also help to discriminate between several forms of biotic homogenisation that took place in different habitats in which the role of Grinnellian and Eltonian filters varies. Accounting for species association has highlighted that differentiated communities, even when composed by habitat specialists, can hide very different processes resulting either in complex and strongly associated networks or very sparse association networks.

## Data availability

All the analyses were conducted using R (version 3.4.4). Data and the R script are available on a Dryad repository (https://doi.org/10.5061/dryad.c2fqz616h) (Rigal *et al*., 2020).

## Notes

### Competing Interest Statement

The authors have declared no competing interest.

