## Supplementary material 1-7 for "Biotic homogenisation in bird communities leads to large-scale changes in species associations": Supplementary_material_GEB20200612.pdf

**Appendix 1**

| **Initial classes** | **Merged classes** | **Main groups** |
|---|---|---|
| Deciduous woodland | Deciduous woodland | Woodland |
| Coniferous woodland | Coniferous woodland | |
| Mixed woodland | Mixed woodland | |
| Young forest | Young forest | |
| Heath | Heath | |
| Scrub | | |
| Others | | |
| Coppice | Coppice | |
| New plantation | | |
| Clear-cut | | |
| Dry limestone meadow | Dry natural meadow | |
| Other dry meadows | | |
| Herbaceous moorland | Moorland | |
| Heather moorland | | |
| Moist natural meadow | Marshland | |
| Flooded meadow | | |
| Reeds | | |
| Salt marsh | | |
| Other open marshland | | |
| Ploughed meadow | Ploughed meadow | Grassland |
| Unploughed meadow | Unploughed meadow | |
| Mixed farmland | Mixed farmland | |
| Open-field | Open-field | |
| Permanent crop | Permanent crop | |
| Other crop | | |
| Urban settlement | Urban settlement | Human settlements |
| Suburban settlement | Suburban settlement | |
| Rural settlement | Rural settlement | |
| Mare | Near open water | |
| Small pond | | |
| Lake | | |
| Reservoir lake | | |
| Waste water | | |
| Stream | | |
| River | | |
| Flooded ditch | | |
| Small canal | | |
| Cliff | Bare rocks | |
| Rocky slope | | |
| Limestone pavement | | |
| Other rocky soil | | |

| |
|---|
| Quarry |
| Mine |
| Dune |

Table S1: Habitat classes summarized from FBBS habitat types.

The poll of 109 species representing 99 % of the total abundance

*Acrocephalus palustris, Acrocephalus schoenobaenus, Acrocephalus scirpaceus, Aegithalos caudatus, Alauda arvensis, Alectoris rufa, Anas platyrhynchos, Anthus pratensis, Anthus trivialis, Apus apus, Ardea cinerea, Buteo buteo, Carduelis carduelis, Chloris chloris, Certhia brachydactyla, Certhia familiaris, Cettia cetti, Cisticola juncidis, Coccothraustes coccothraustes, Coloeus monedula, Columba livia, Columba oenas, Columba palumbus, Corvus corone, Corvus frugilegus, Coturnix coturnix, Cuculus canorus, Cyanistes caeruleus, Cygnus olor, Delichon urbicum, Dendrocopos major, Dendrocoptes medius, Dryobates minor, Dryocopus martius, Emberiza calandra, Emberiza cia, Emberiza cirlus, Emberiza citrinella, Emberiza schoeniclus, Erithacus rubecula, Falco tinnunculus, Fringilla coelebs, Fulica atra, Galerida cristata, Gallinula chloropus, Garrulus glandarius, Hippolais polyglotta, Hirundo rustica, Jynx torquilla, Lanius collurio, Larus argentatus, Larus ridibundus, Linaria cannabina, Locustella naevia, Lophophanes cristatus, Loxia curvirostra, Lullula arborea, Luscinia megarhynchos, Merops apiaster, Milvus migrans, Motacilla alba, Motacilla cinerea, Motacilla flava, Muscicapa striata, Oenanthe oenanthe, Oriolus oriolus, Perdix perdix, Periparus ater, Parus major, Passer domesticus, Passer montanus, Phasianus colchicus, Phoenicurus ochruros, Phoenicurus phoenicurus, Phylloscopus bonelli, Phylloscopus collybita, Phylloscopus sibilatrix, Phylloscopus trochilus, Pica pica, Picus viridis, Podiceps cristatus, Poecile montanus, Poecile palustris, Prunella modularis, Pyrrhula pyrrhula, Regulus ignicapilla, Regulus regulus, Saxicola rubetra, Saxicola torquatus, Serinus serinus, Sitta europaea, Streptopelia decaocto, Streptopelia turtur, Sturnus vulgaris, Sylvia atricapilla, Sylvia borin, Sylvia cantillans, Sylvia communis, Sylvia curruca, Sylvia melanocephala, Sylvia undata, Tadorna tadorna, Turdus pilaris, Troglodytes troglodytes, Turdus merula, Turdus philomelos, Turdus viscivorus, Upupa epops, Vanellus vanellus.*

**Appendix 2**

Clique structure

Many alternative network indices are available to study the complexity or structure of a network (May 1974, Landi et al. 2018, Mariani et al. 2019). For instance, the degree distribution or the connectance respectively reveal densely associated species or networks with a high density of associations. Moreover, motifs defined as recurring and significant patterns of interconnections (Milo et al. 2002, Shen-Orr et al. 2002) can reveal different network classes. Here, we chose to focus on cliques to assess whether networks were structured as tightly knit sets of species (Pattillo et al. 2013). We thus extended the concept of transitivity (or global clustering coefficient), corresponding to the ratio between the number of observed 3-cliques and the maximum possible number of 3-cliques (Watts & Strogatz 1998, Barrat & Weigt 2000), by using cliques with three *and more* species.

Network examples

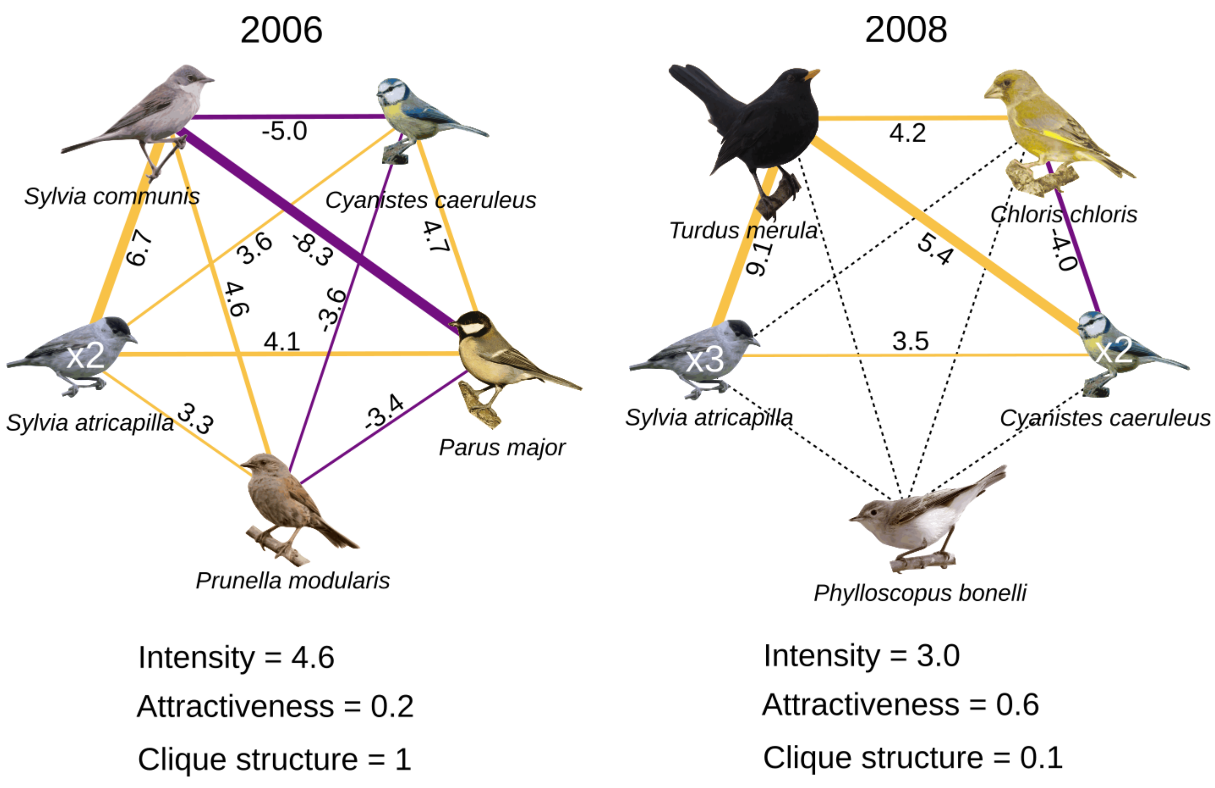

Figure S2: Species association networks in 2006 and in 2008 from a sampling point located in an open-field area in the Atlantic region. Three species (*Sylvia communis*, *Prunella modularis* and *Parus major*) initially observed in 2006 were not seen in 2008. Three new species were observed in 2008: *Turdus merula*, *Chloris chloris* and *Phylloscopus bonelli*. Positive associations are in light orange and negative associations in dark purple. Non-significant associations are represented by dotted lines. Pairwise association values are given, and for each species, the number of individuals is specified when above 1. Between 2006 and 2008, the average association strength and the clique structure decreased while the number of positive associations relatively increased.

**Appendix 3**

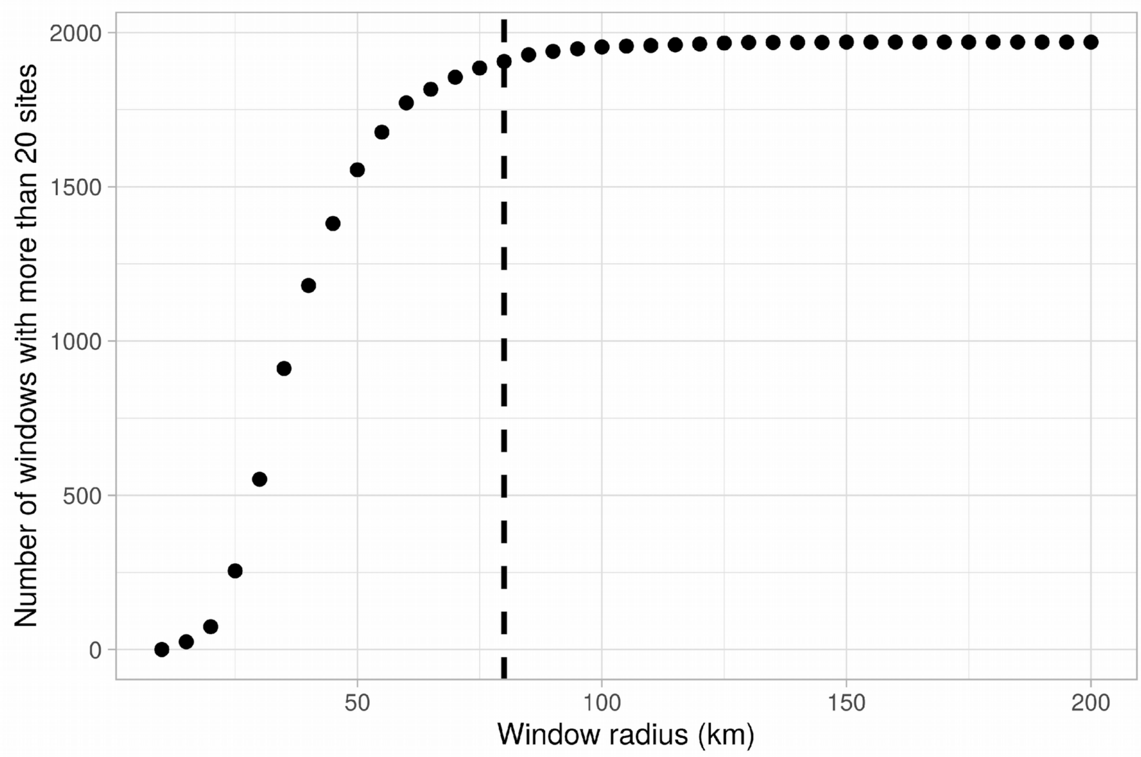

Figure S3a: Number of windows with at least 20 sites *vs.* window radius. Dashed line shows the selected radius (80 km).

Spatial and temporal variation in network indices and β-diversity

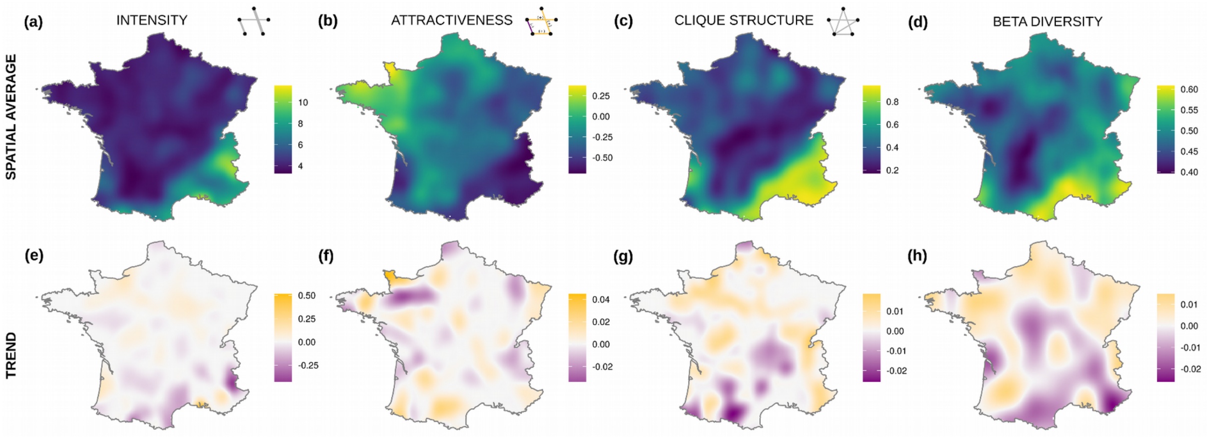

Figure S3b: Spatial distribution of three indices of the association networks and β-diversity of

bird communities. First row: Average spatial values (2001 to 2017) of (a) intensity, (b) attractiveness, (c) clique structure and (d) β-diversity. Low values are represented in dark blue and high values in light yellow. Second row: spatial distribution of the temporal trends of (e) intensity, (f) attractiveness, (g) clique structure of the association networks between 2001 and 2017 and (h) trend of β-diversity in bird communities between 2001 and 2017. Negative values are represented in dark purple, null in grey and positive in light orange.

Intensity (*i.e.* the mean association strength in the network) was high in Mediterranean (south-eastern) (Fig. S3b). Attractiveness (*i.e.* the ratio of positive versus negative associations) was negative in most eastern France. Clique structure (*i.e.* the ratio of existing versus possible cliques) was low in most parts of France except for the Mediterranean and Alpine (south-eastern) areas.

The temporal trend in intensity increased in the northern areas and decreased in the southern areas. Temporal changes in attractiveness were more heterogeneously distributed. The clique structure decreased in south-central and south-western France but increased elsewhere.

β-diversity showed higher values in the Mediterranean region (south-eastern), and more generally in eastern France, than in other regions. Spatial distribution of trends in β-diversity shows an important decrease in southern areas and weak increase in northern and western areas.

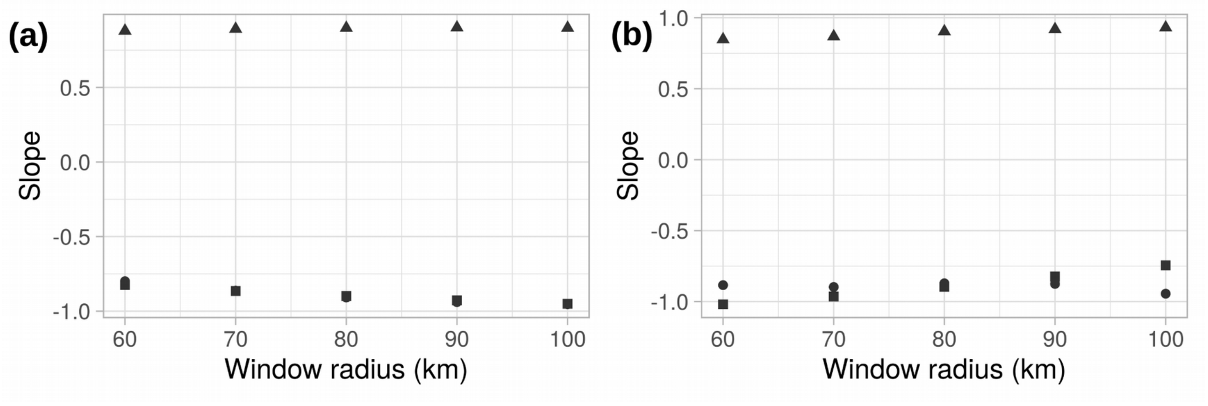

Figure S3c: Linear regression coefficient between window indices calculated at the local (i.e. the site) scale (window radius = 0) and for each window size in space (a) and for index trend (b). Linear regression coefficient between intensity and clique structure (▲), intensity and attractiveness (●) and attractiveness and clique structure (■).

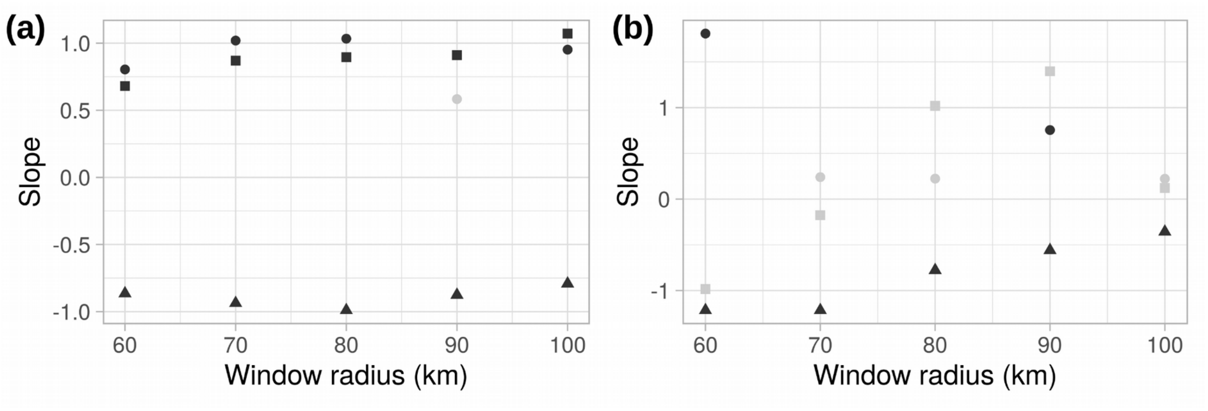

Figure S3d: Linear regression coefficients between indices and β-diversity (a) and between their trends (b) calculated for each window size. Linear regression coefficient between β-diversity and attractiveness (▲), intensity (●) and clique structure (■). Non significant values are in grey.

We found similar relationships between network indices for window size ranging from 60 km to 100 km radius (Fig. S3c). The positive relationship between intensity and clique structure remained constant. The relationship between intensity and attractiveness stayed negative as well as the relationship between clique structure and attractiveness. Similar relationships were found using temporal trends.

We found similar relationships between network indices and β-diversity for window size ranging from 60 km to 100 km radius (Fig. S3d). The positive relationship between intensity and β-diversity remained constant (non significant only for a window size of 90 km). The relationship between attractiveness and β-diversity stayed negative and the relationship between clique structure and β-diversity remains positive. A negative relationship was also found between attractiveness and β-diversity using temporal trends. And to the exception of a window size of 90 km, temporal relationships between intensity and β-diversity or between clique structure and β-diversity stay non significant.

**Appendix 4**

β-diversity, species richness, network indices

To test whether network indices and β-diversity were intrinsically linked, *i.e.* numerically redundant, we used two approaches to measure their redundancy: a simulation approach and a permutation approach.

Simulation

The simulation approach allows to disentangle the potential redundancy that could have emerged from co-occurrence patterns. If some species co-occur little with others this could be reflected both in the computation of associations (negative associations) and in the originality of the community to which they belong (leading to high β-diversity). This relationship between β-diversity and associations due to co-occurrence patterns could impact the link between β-diversity and network indices. In such a case, β-diversity and network indices would be redundant. If this is the case, however, the relationship should be also observed in simulated data.

We therefore used simulated data mimicking species co-occurrence patterns and calculated associations, network indices and β-diversity on those simulated data. The simulated dataset was composed of the abundance of 15 species in 400 sites (10 sampling points per site). Each site was monitored for 5 years, randomly drawn between 2001 and 2017. Only one habitat and one biogeographic region were considered. Sites were located on a square grid (285 x 285 km), centred on France, and equally separated by 15 km. Associations between species were set as in Figure S4a to aggregate and segregate species and association values were then obtained through the framework used in the main text. Network indices and β-diversity were then calculated from these simulated data and compared using the same model as described in the main text.

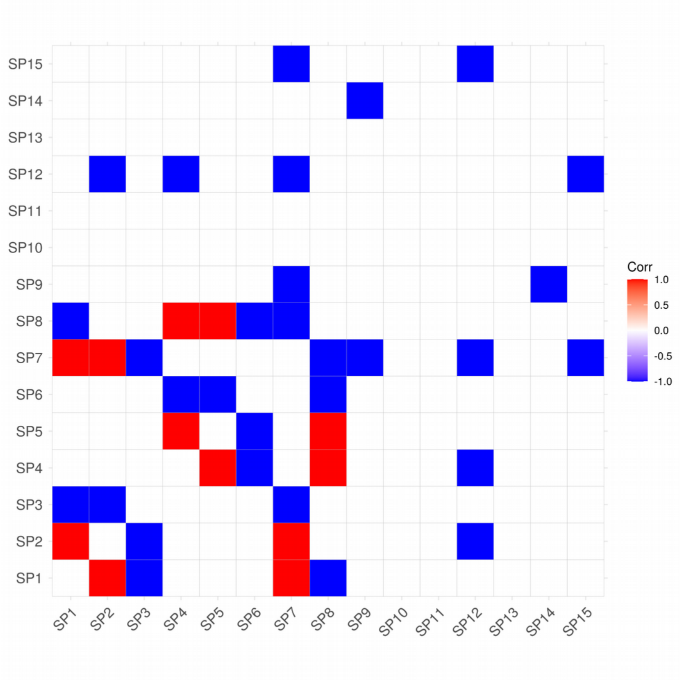

Figure S4a: Simulated associations between simulated species. Negative associations are shown in blue and positive associations in red.

Overall, we found that simulated network indices were not related to β-diversity neither in space nor in time (Fig. S4b).

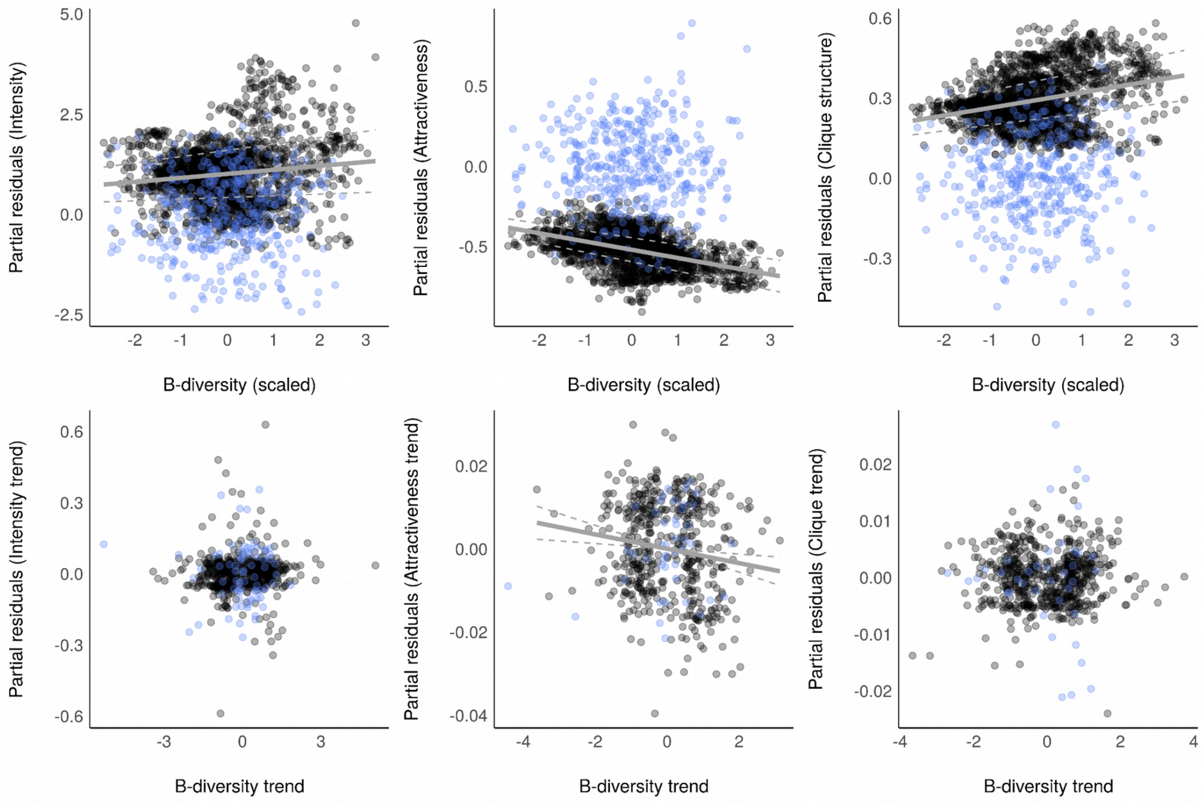

Figure S4b: Relationship between network indices computed on simulated (blue dots) versus original data (black dots). Only significant relationships are shown by solid lines with confidence intervals in dashed lines.

### Permutation

We used a permutation approach to complete the simulation approach. Simulated data were constructed to mimic the original data but they could still be considered as simplified compared to the original data, whereas permuted data have the same characteristic as the original data. To obtain permuted data, we reshuffled species identity between all sampling points, keeping constant the number of species in each sampling point. We used significant species associations computed on the original data to reconstruct permuted species association networks. We calculated network indices and β-diversity on the permuted communities and we analysed their relationship using the same model as described in the main text.

We found that β-diversity values for permuted data were lower than original data, as expected due to the permutation. Permuted network indices were related to β-diversity in an opposite way compared to the relation obtained with original data (Fig. S4c), with the exception of attractiveness in time.

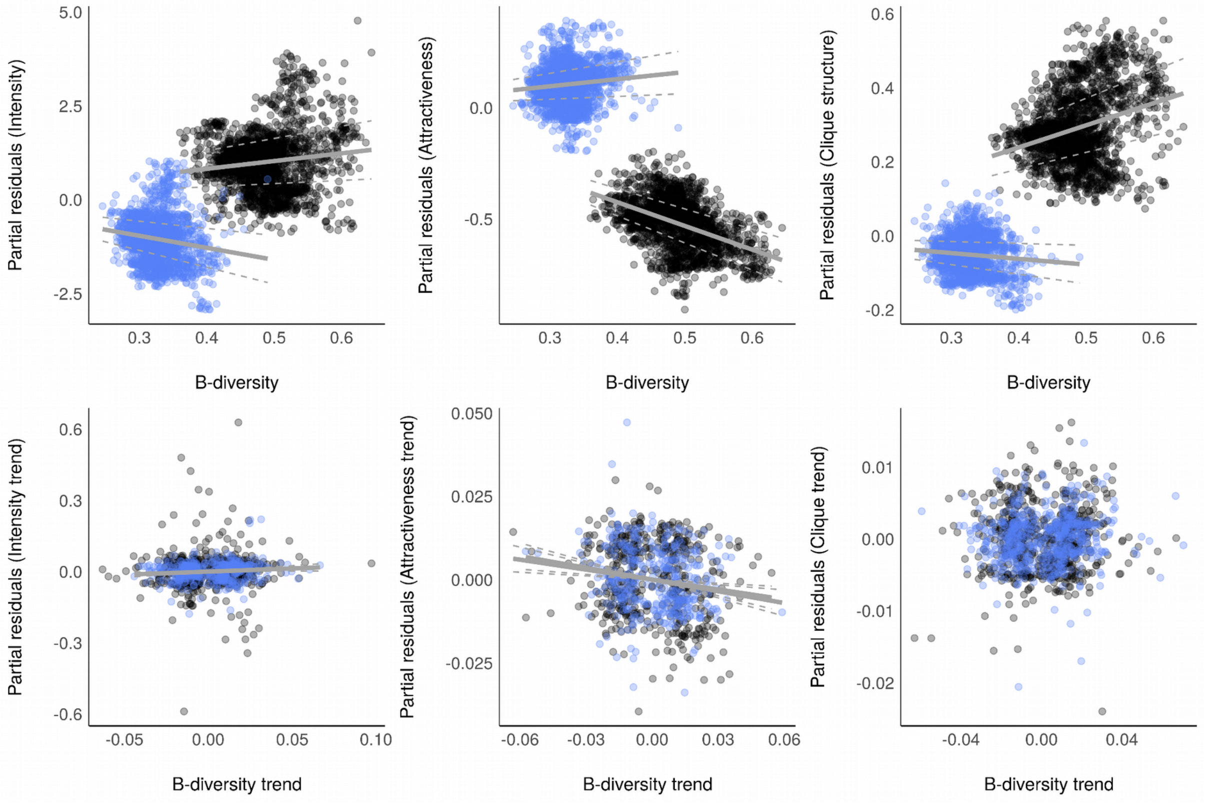

Figure S4c: Relationship between network indices computed on permuted (blue dots) versus original data (black dots). Only significant relationships are shown by solid lines with confidence intervals in dashed lines.

**Appendix 6**

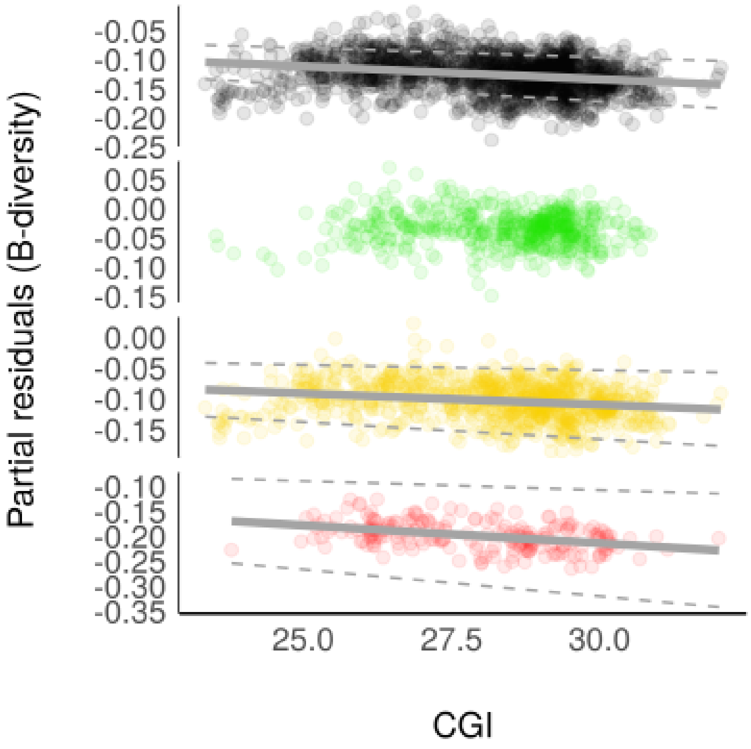

Figure S6a: Relationship between β-diversity and CGI in space, for all habitats (black dots), woodland (green dots), grassland (yellow dots) and human settlements (red dots). Dots correspond to partial residuals of the regression models regressed over predictors and regression lines (solid lines) with confidence intervals (dashed lines) are shown when significant.

Overall, controlling for the species richness, we found that β-diversity was negatively related to CGI in space (slope = $-4.10^{-3}$, sd = $1.10^{-3}$, t-value = -7, p-value < $1.10^{-10}$, df = 1943, adjusted $r^2 = 0.63$) and time (slope = -0.03, sd = 0.01, t-value = -5, p-value = $6.10^{-7}$, df = 1296, adjusted $r^2 = 0.06$) (Fig. S6a), meaning that the more original species turnover is, the more communities are composed of habitat specialists. In other words, the more biotic homogenisation (low β-diversity), the more habitat generalist communities. This, however, was not verified in forest areas where β-diversity was not related to the relative number of generalists in communities (slope = $-1.10^{-3}$, sd = $1.10^{-3}$, t-value = -1, p-value = 0.31, df = 548, adjusted $r^2$ = 0.51), contrary to grassland (slope = $-4.10^{-3}$, sd = $1.10^{-3}$, t-value = -4, p-value = $9.10^{-5}$, df = 987, adjusted $r^2$ = 0.61) or human settlements (slope = $-7.10^{-3}$, sd = $2.10^{-3}$, t-value = -4, p-value = $8.10^{-5}$, df = 221, adjusted $r^2$ = 0.72).

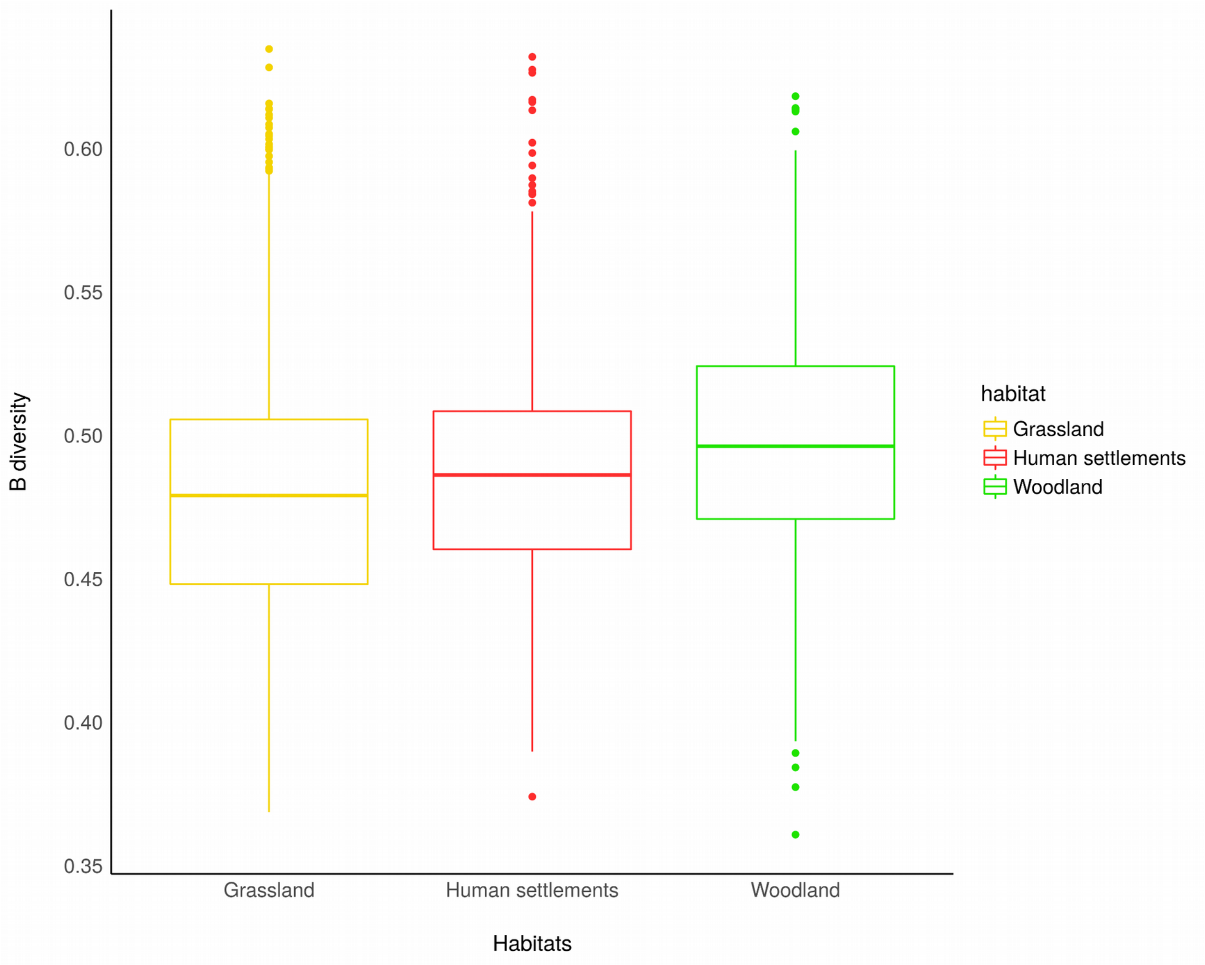

Figure S6b: β-diversity across the three main habitats.

Controlling for the species richness, β-diversity was found higher in woodland (Fig. S6b) than in other habitats (reference: Grassland, estimate = $6.9.10^{-3}$, sd = $1.6.10^{-3}$, t = 4, p = $2.10^{-5}$). β-diversity in human settlement was intermediate between woodland and grassland (reference: Grassland, estimate = $6.7.10^{-3}$, sd = $2.2.10^{-3}$, t = 3, p = $2.4.10^{-3}$).

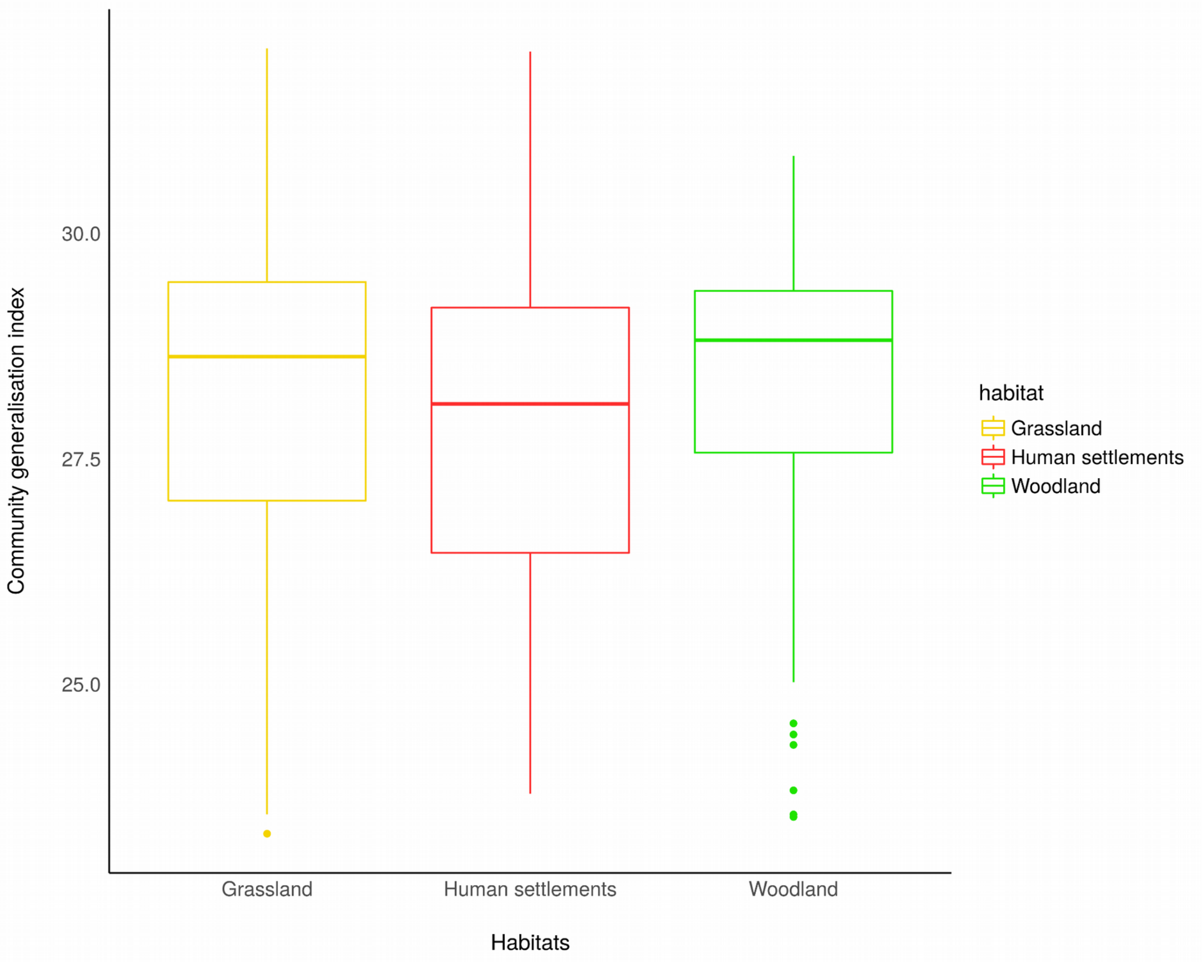

Figure S6c: CGI for each of the three main habitats.

Controlling for the species richness, CGI was found higher in woodland (Fig. S6c) than in other habitats (reference: Grassland, estimate = 0.24, sd = 0.06, t = 4, p = $3.10^{-5}$). CGI in human settlement was lower than in other habitats (reference: Grassland, estimate = -0.24, sd = 0.08, t = -3, p = $2.10^{-3}$).

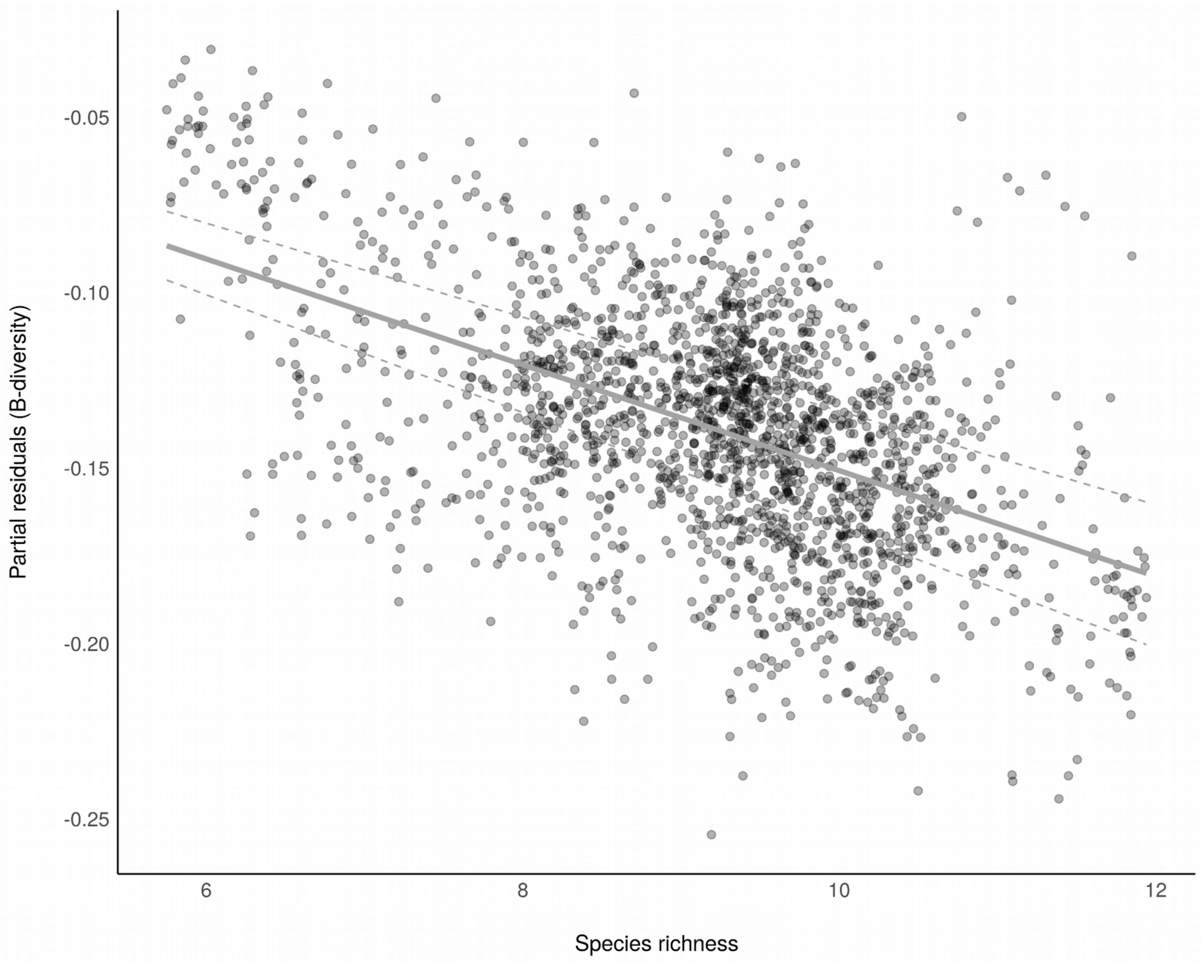

Figure S6d: Relationship between β-diversity and species richness. Dots correspond to partial residuals of the GAMM. The regression line is shown by the solid line with confidence intervals as dashed lines.

β-diversity calculated as the species turnover component and thus independent from the nestedness component was still found negatively related to the species richness (Fig. S6d) (slope = -0.015, sd = 0.001, t-value = -17, p-value < $1.10^{-10}$, df = 1943, adjusted $r^2$ = 0.62).

**Appendix 7**

Species functional dissimilarity

Functional dissimilarity was estimated using a life-history trait dataset for the birds of the Palaearctic (Storchová & Hořák 2018). Life history traits are composed of functional traits (lengths of bird, wing, tail, bill, tarsus, weight, sexual dimorphism, clutch size, broods per year, egg length, width and weight, type of young, association during nesting, nest type, nest building, mating system, incubation period, incubation sex, territoriality, migration strategy (sedentary, facultative, short or long distance migrant)), diet (at least 10% of this diet during the breeding season, folivore, frugivore, granivore, arthropods, other invertebrates, fish, other vertebrates, carrion, omnivore) and habitat requirements (breeding area with deciduous forest, coniferous forest, woodland, shrub, savanna, tundra, grassland, mountain meadows, reed, swamps, desert, freswater, marine, rocks, human settlements).

We computed the Generalized Functional Diversity (GFD) following Mouchet et al. (2008) to generate a pairwise functional dissimilarity distance between all pairs of species (Mouchet et al. 2010). This method computes all the combinations of the clustering algorithms from a functional distance matrix and provides the best consensus tree, *i.e.* the one for which the dissimilarity with the original distance matrix is the lowest according to the 2-norm goodness-of-fit index (Mérigot et al. 2010). We constructed our initial functional distance matrix using the Gower’s distance because the trait matrix contained both qualitative and quantitative data (Gower 1971). For our data, the best consensus dendrogram was built using the Unweighted Pair Group Method using arithmetic Average algorithm (UPGMA) (2-norm=3.39, threshold=3.47). From this dendrogram, we computed the cophenetic functional distance (Sokal & Rohlf 1962) between species, which is hereafter called “pairwise functional dissimilarity”.

Community functional diversity

We calculated the average functional diversity in each community (CFD) using a community weighted mean of the functional dissimilarity between species pairs (obtained as explained above). We then analyse the difference in CFD between the three main habitats (woodland (coniferous, deciduous or mixed forest), grassland (ploughed and unploughed meadows, mixed farmland, open-field or crop) and human settlements (urban settlements, suburban settlements, rural settlements)).

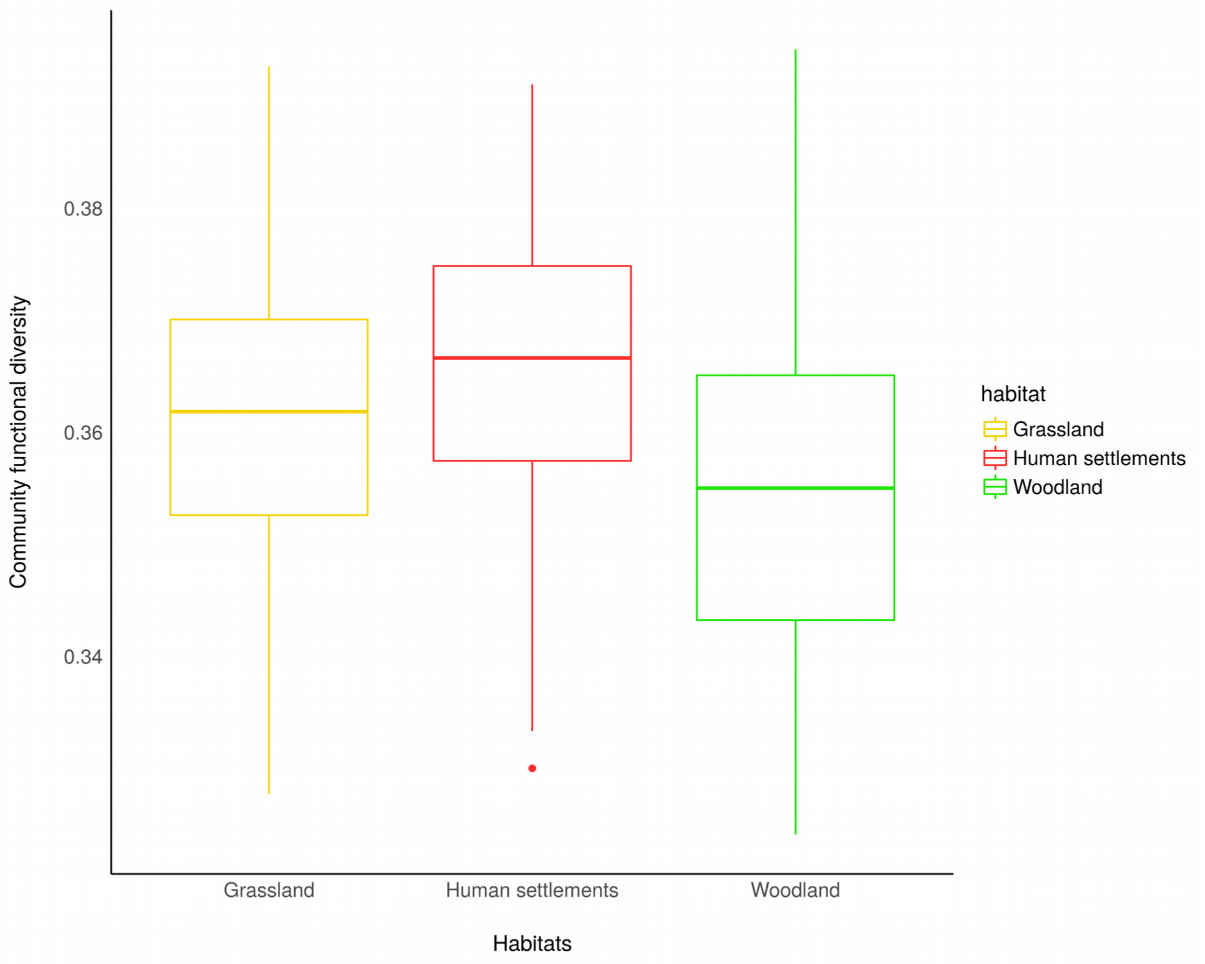

Figure S7: Community functional diversity (CFD) in each of the three main habitats, showing the highest in human settlements and the lowest in woodland.

Overall, controlling for the species richness, CFD was found higher in human settlement than in other habitats (Fig. S7) (reference: Grassland, estimate = $4.3.10^{-3}$, sd = $0.8.10^{-3}$, t = 6, p = $1.10^{-8}$). CFD was lower in woodland than in other habitats (reference: Grassland, estimate = $-3.7.10^{-3}$, sd = $0.6.10^{-3}$, t = -7, p < $1.10^{-10}$).
